# The regulation of liver gene expression by carbohydrates is mouse strain specific

**DOI:** 10.1101/2020.11.11.378497

**Authors:** Yuling Chi, Dou Yeon Youn, Alus M. Xiaoli, Li Liu, Jacob B. Pessin, Fajun Yang, Jeffrey E. Pessin

## Abstract

C57BL/6J and BALB/cJ mouse strains were analyzed by deep mRNA sequencing of the liver in the fasted state and following ingestion of standard laboratory mouse chow supplemented with plain drinking water or water containing 20% glucose, sucrose or fructose. Supplementation with these carbohydrates induced unique extents and temporal changes in gene expressions in a strain specific manner. Fructose and sucrose stimulated gene changes peaked at 3 h postprandial, whereas glucose effects peaked at 12 h postprandial in C57BL/6J mice and at 6 h postprandial in BABL/cJ mice. Network analyses revealed that fructose changed genes were primarily involved in lipid metabolism and were more complex in C57BL/6J than in BALB/cJ mice. These data demonstrate that there are qualitative and quantitative differences in the normal physiological responses of the liver between these two strains of mice and C57BL/6J is more sensitive to sugar intake than BALB/cJ.

## Introduction

Carbohydrates, naturally existing in grains, fruits and other food sources, are major components of our diets. Extra carbohydrates, for instance table sugar, are often added to our food and beverages, and the amount of sugar-like sweeteners supplemented into food and beverages has been increasing [1]. Excessive consumptions of sugar have been shown directly or indirectly to associate with metabolic disorders including obesity and diabetes [2–5]. Excessive consumptions of sugars, including disaccharide, such as sucrose, and monosaccharides, such as glucose and fructose, to different extents result in abnormalities in metabolic processes such as gluconeogenesis and lipogenesis in the liver. For instance fructose leads to liver lipid accumulation, dyslipidemia and increased uric acid levels [2]. However, the molecular mechanisms and signaling pathways of how these carbohydrates affect metabolic processes are not completely understood and remain somewhat controversial [2].

Analyses of high carbohydrate diets on genome wide gene expressions in animal models have provided valuable information on gene changes in response to excessive sugar intake, important for better understanding mechanisms at the molecular level [6–11]. Yet, the designs and/or the technologies used in most of reported investigations are limited. In some studies glucose, sucrose or fructose was given to mice in combination with high fat diet. In most studies only the nutrient-sensitive C57BL/6J mouse was used. Most studies were investigating only the long-term (several weeks to several months) effects. There has been no comprehensive study on the acute effects of individual sugars on the genome wide gene expressions.

Recently we reported temporal and dynamic genome-wide gene expression changes in livers of nutrient-sensitive C57BL/6J and nutrient-insensitive BALB/cJ mice from fasted to fed states, revealed by utilizing deep mRNA-seq technology [12]. We have also included separate groups of these two strains of mice given acute supplementation with one of three carbohydrates (glucose, sucrose and fructose). Here we report the impact of supplementing low fat chow diets with three individual carbohydrates on gene expression changes in liver, and its implications in metabolic processes and signaling networks and pathways.

## Materials and Methods

### Mice

All following experimental procedures performed with mice were approved by the Institutional Care and Use Committee at the Albert Einstein College of Medicine in accordance with the “Guide for the Care and Use of Laboratory Animals” published by the National Institute of Health. Wild type C57BL/6J and BALB/cJ mice at age of 8 weeks were purchased from the Jackson Laboratory. One day after arrival, they were trained for 1 week for regulated fasting (from 10 pm to 7 pm) and feeding of low-fat chow diet (PicoLab Mouse Diet 5053, 10 % of calories from fat and 64.5% of calories from carbohydrates) (from 7 pm to 10 pm). After one week of training, mice were divided into four groups. All of them were fasted from 10 pm the day before to 7 pm the day of the experiment. Water was taken away from them from 4 pm to 7 pm only on the experiment day. At 7 pm, they were provided with food and water supplemented with or without 20% (g/g) of glucose or sucrose or fructose. The mice were allowed to eat *ad libitum* until 10 pm at which point food was removed and water was changed to regular water without any supplement. Mice were sacrificed at 7 pm as fasted controls and at 10 pm (3 h postprandial), 1 am (6 h postprandial) and 7 am (12 h postprandial) following the initiation of feeding, respectively. Livers were harvested and snap frozen in liquid nitrogen. This protocol resulted in the generation of 26 groups of mice with 4-5 replicate mice per group (Table 1).

**Table 1.** Information on groups of samples.

| Group | Strain | Treatment | Time following feeding (hour) | n | Group | Strain | Treatment | Time following feeding (hour) | n |
| --- | --- | --- | --- | --- | --- | --- | --- | --- | --- |
| **g1** | C57BL/6/J | Fasted | 0 | 5 | **g2** | BALB/cJ | Fasted | 0 | 5 |
| **g3** | C57BL/6/J | Fed Control | 3 | 5 | **g7** | BALB/cJ | Fed Control | 3 | 5 |
| **g4** | C57BL/6/J | Fed Glucose | 3 | 5 | **g8** | BALB/cJ | Fed Glucose | 3 | 5 |
| **g5** | C57BL/6/J | Fed Sucrose | 3 | 5 | **g9** | BALB/cJ | Fed Sucrose | 3 | 5 |
| **g6** | C57BL/6/J | Fed Fructose | 3 | 5 | **g10** | BALB/cJ | Fed Fructose | 3 | 5 |
| **g11** | C57BL/6/J | Fed Control | 6 | 5 | **g15** | BALB/cJ | Fed Control | 6 | 5 |
| **g12** | C57BL/6/J | Fed Glucose | 6 | 5 | **g16** | BALB/cJ | Fed Glucose | 6 | 5 |
| **g13** | C57BL/6/J | Fed Sucrose | 6 | 5 | **g17** | BALB/cJ | Fed Sucrose | 6 | 5 |
| **g14** | C57BL/6/J | Fed Fructose | 6 | 5 | **g18** | BALB/cJ | Fed Fructose | 6 | 5 |
| **g19** | C57BL/6/J | Fed Control | 12 | 5 | **g23** | BALB/cJ | Fed Control | 12 | 5 |
| **g20** | C57BL/6/J | Fed Glucose | 12 | 5 | **g24** | BALB/cJ | Fed Glucose | 12 | 4 |
| **g21** | C57BL/6/J | Fed Sucrose | 12 | 5 | **g25** | BALB/cJ | Fed Sucrose | 12 | 5 |
| **g22** | C57BL/6/J | Fed Fructose | 12 | 5 | **g26** | BALB/cJ | Fed Fructose | 12 | 4 |

### Total RNA extraction

Approximately 5mg of frozen liver powder was completely dissolved in 500 μL of Trizol reagent at room temperature. One hundred μL of chloroform was added to the Trizol-liver mixture. The mixture was kept at room temperature for 2 min and subsequently centrifuged at 12,000 rpm for 15 min at 4°C. The top layer (aqueous) was isolated and mixed with 100% ethanol at 1:1 ratio. Up to 700 μL of the mixture was transferred onto RNeasy mini column from Qiagen kit (catalogue No. 74106). Thereafter the rest of procedure of the same protocol part 1 from step 3 to the end was followed to finish total RNA extraction. All total RNA samples were sent to Novogene Corporation Inc in Sacramento, CA 95826 for genome wide mRNA sequencing. All samples passed through the following three steps before library construction: 1) Nanodrop for RNA purity check, OD260/OD280 in a range of 1.95 – 2.05; 2) agarose gel electrophoresis for RNA integrity and potential contamination; and 3) Agilent 2100 for confirming RNA integrity.

### Library construction and sequencing

Library construction and sequencing were conducted by Novogene. Briefly, mRNA was purified from total RNA by using poly-T oligo attached magnetic beads and fragmented. cDNA was synthesized. cDNA fragments with 150 bp, paired and two-ended reads were generated and selected. Libraries (30,000,000 fragments / library) were constructed and fed into Illumina machines.

### Data Processing

STAR v2.5 was used to align clean data to mm10 reference genome with parameter mismatch = 2 [13]. HTSeq v 0.6.1 was used to count the read numbers mapped to each gene [14]. Differential expression analysis between two conditions/groups was performed using the DESeq2 R package (v 2_1.6.3), which generated log2 fold change and adjusted *p* value (p_adj_) using Benjamini-Hochberg method [15].

Fragments per kilobase of transcript per million mapped reads (FPKMs) were calculated to normalize read counts. Based on FPKM of each gene of each sample, square of Pearson coefficients (R^2^s) were calculated to show correlations between samples and reproducibility. FPKM of each gene was averaged in each group, and log10 (FPKM+1) values were used to generate a heatmap, in which groups and genes were clustered, respectively. Euclidean distance between groups were also calculated based on log10 (FPKM+1) of all genes. Differentially expressed genes were annotated and classified with Gene Ontology terms using Panther Classification system [15], and networks were generated using Ingenuity Pathway Analysis (IPA) [16].

## Results

### Data validation

As shown in Table 1, in this study, there were 26 groups with 5 mouse replicates for each group except groups 24 and 26, which had 4 replicates, resulting in a total of 128 liver samples from 128 individual mice. The RNA-seq data for all samples have been deposited in the GEO database with accession number GSE 137385. The quality of mRNA-seq data from the 128 samples is summarized in supplementary S1 Table, with error rates for all samples < 0.03% and > 90% of reads with Q30. Correlation coefficients (R^2^s) of gene expression reads as FPKM of all samples within each group are listed in S2 Table. All R^2^s are greater than 0.92. The correlation coefficient matrix of all samples shown in S1 Fig demonstrates clear patterns, confirming that all the data within each group were well correlated. Zoomed in correlation plots between any two samples in group 1 are shown in our recently published report [12]. All of these data indicate that all the mRNA-seq results were reproducible and reliable.

### Genome wide gene expression and clustering analysis

We then carried out hierarchical clustering analysis of gene expressions in all 26 groups. Average FPKM for each gene of each group was calculated and log10 (FPKM+1) of all genes were plotted in a heatmap (Fig 1A). This heatmap shows clear separation between two strains of mice. At various time points, intra-strain groups closely clustered while inter-strain groups separated from each other. The 6 hour postprandial groups of C57BL/6J mice are furthest away from those groups of BALB/cJ mice. Eucledean distances among groups were calculated based on log10 (FPKM+1) of all genes and are shown in Fig 1B. The arbitrary distance values are shown in Fig 1C. These data demonstrate that there are apparent differences in hepatic gene expressions between C57BL/6J and BALB/cJ mice and the greatest differences occurred at 6 hour postprandial, although the overall distances at 6 hour are only slightly bigger than that at 3 hour (Fig 1C).

**Fig 1.**
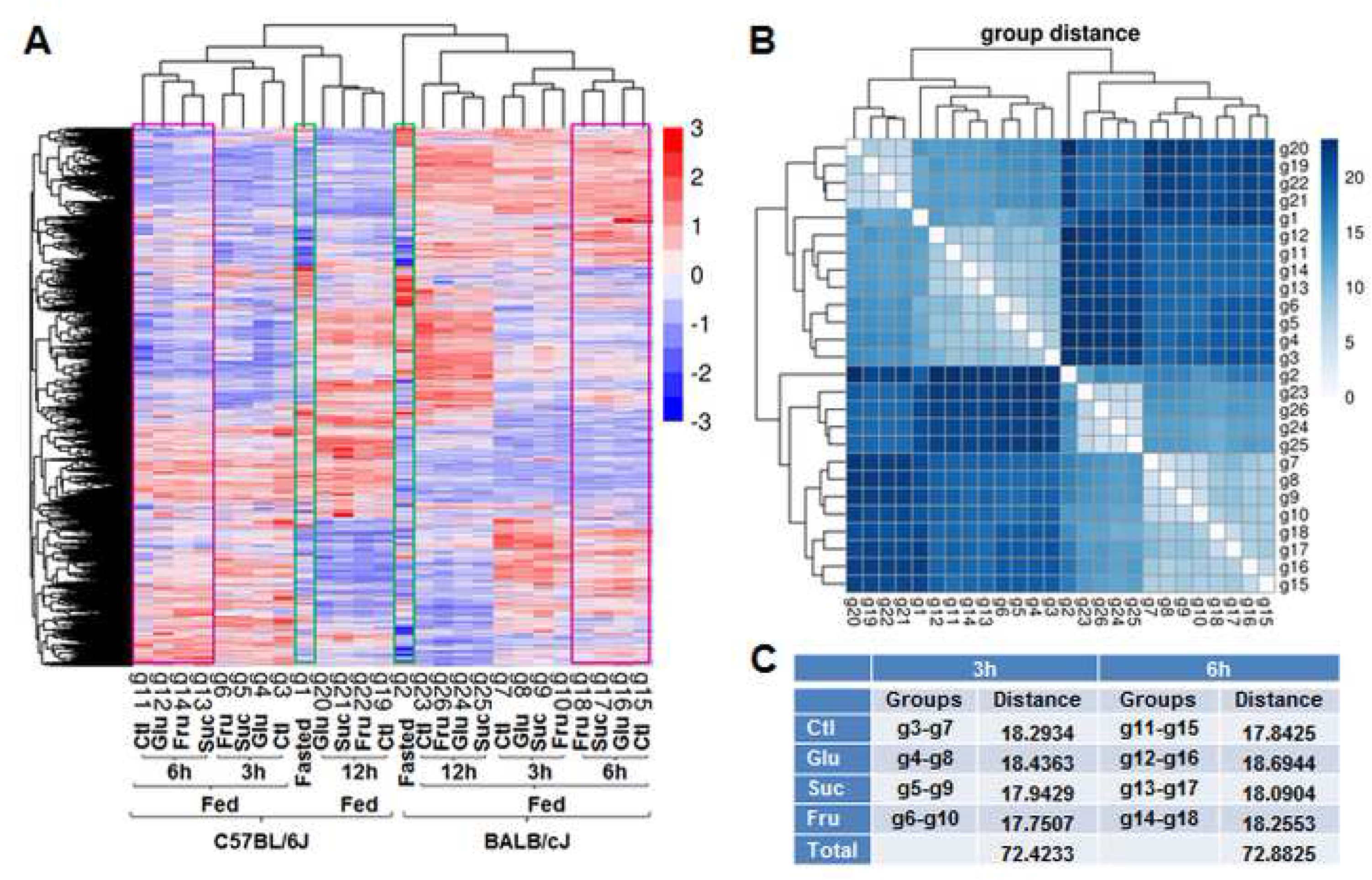
Genome wide analyses of expressed genes in two strains of mice. (A) Cluster analysis of expressed genes in 26 groups. Hierarchical clustering analysis was carried out with log10(FPKM+1) of expressed genes of all 26 groups under different experimental conditions as indicated. (B) Euclidean distance among those 26 groups based on log10(FPKM+1). (C) Arbitrary values of distances.

### Differentially expressed genes in two strains of mice

To investigate the gene expression differences between C57BL/6J and BALB/cJ mice in details, we directly compared gene expressions in those two strains. Gene populations of two strains are shown in the Venn diagrams (Fig 2). Here, average FPKM of each gene in the same group was used. If average FPKM ≥ 1, we viewed this gene as expressed in the group. Otherwise, we viewed it as unexpressed in the group. Under all conditions shown in Fig 2, the numbers of shared genes by both strains were in the range of 10,139 to 10,471. The numbers of genes uniquely expressed in C57BL/6J mice were in the range of 319 to 531, whereas the numbers of unique genes expressed in BALB/cJ mice were in the range of 403 to 602, slightly higher than that in C57BL/6J mice.

**Fig 2.**
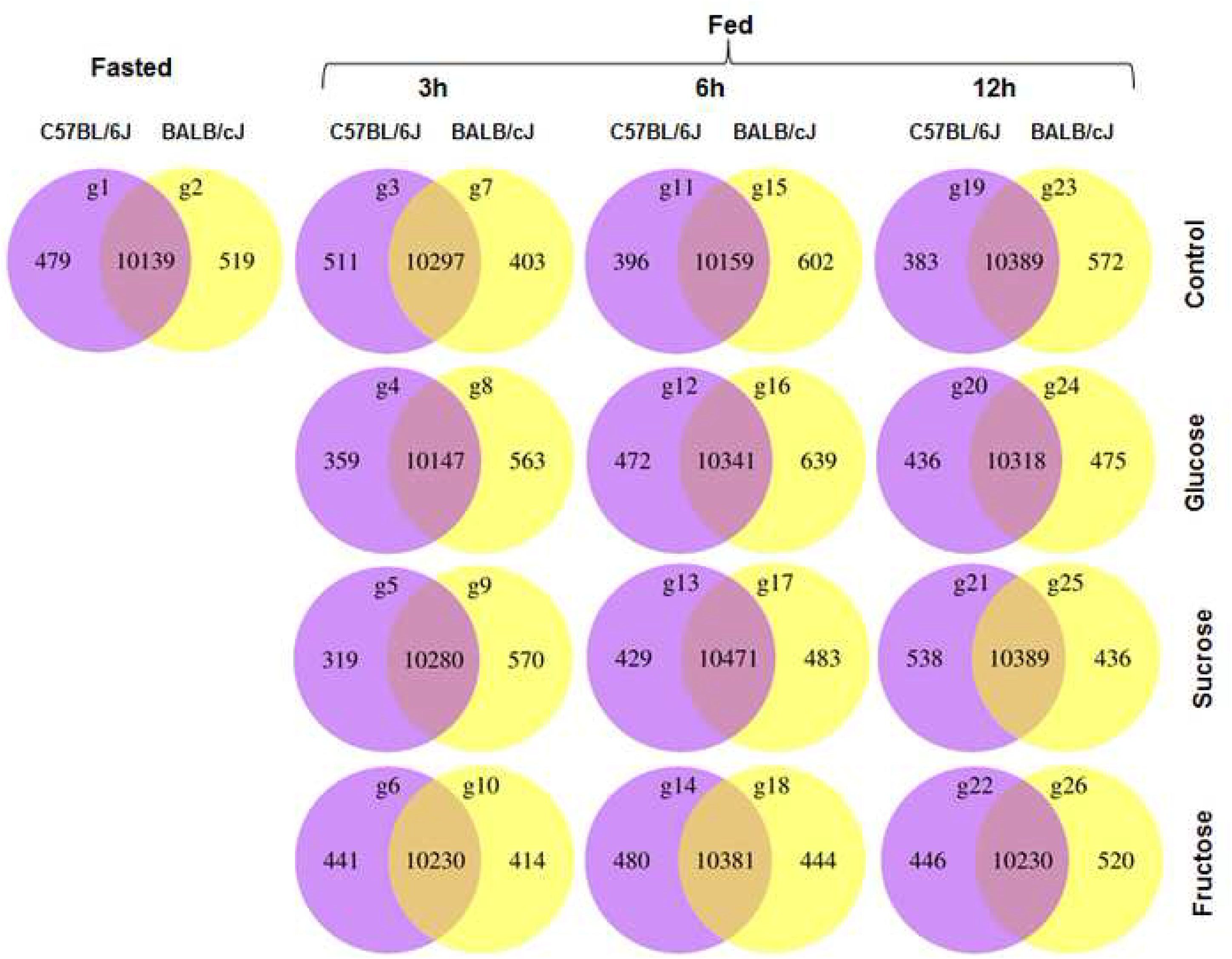
Gene distributions and populations in C57BL/6J and BALB/cJ mice in the fasted state and fed without or with different carbohydrate supplements. Genes expressed in C57BL/6J mice are in purple and genes expressed in BALB/cJ mice are in yellow. Genes with FPKM ≥ 1 were considered expressed. Genes with FPKM < 1 were considered absent.

To better understand the functions of those uniquely expressed genes in each strain under different conditions, we classified genes using Panther Classification program. Detailed analyses and classifications of genes uniquely expressed in C57BL/6J or BALB/cJ mice in the fasted state and at 6 hour postprandial without carbohydrate supplement have been reported in our recent publication [12]. In this report, we focus on the gene expressions with carbohydrate supplements. At 6 hour postprandial time point, the numbers of genes uniquely expressed in C57BL/6J mice under the condition of without carbohydrate (control), or with glucose, sucrose, or fructose supplement are 396, 472, 429 and 480 respectively. The numbers of genes uniquely expressed in BALB/cJ mice under the condition of without carbohydrate (control), or with glucose, sucrose, or fructose supplement are 602, 639, 483 and 444 respectively. We uploaded these 8 sets of genes onto Panther Classification program and extracted the genes involved in metabolic processes class (S3 Table). Further classification of genes in S3 Table resulted in a list of genes participating in the metabolisms of three major types of macronutrients, *i.e*. lipids, carbohydrates, and proteins/amino acids (Table 2).

**Table 2.**
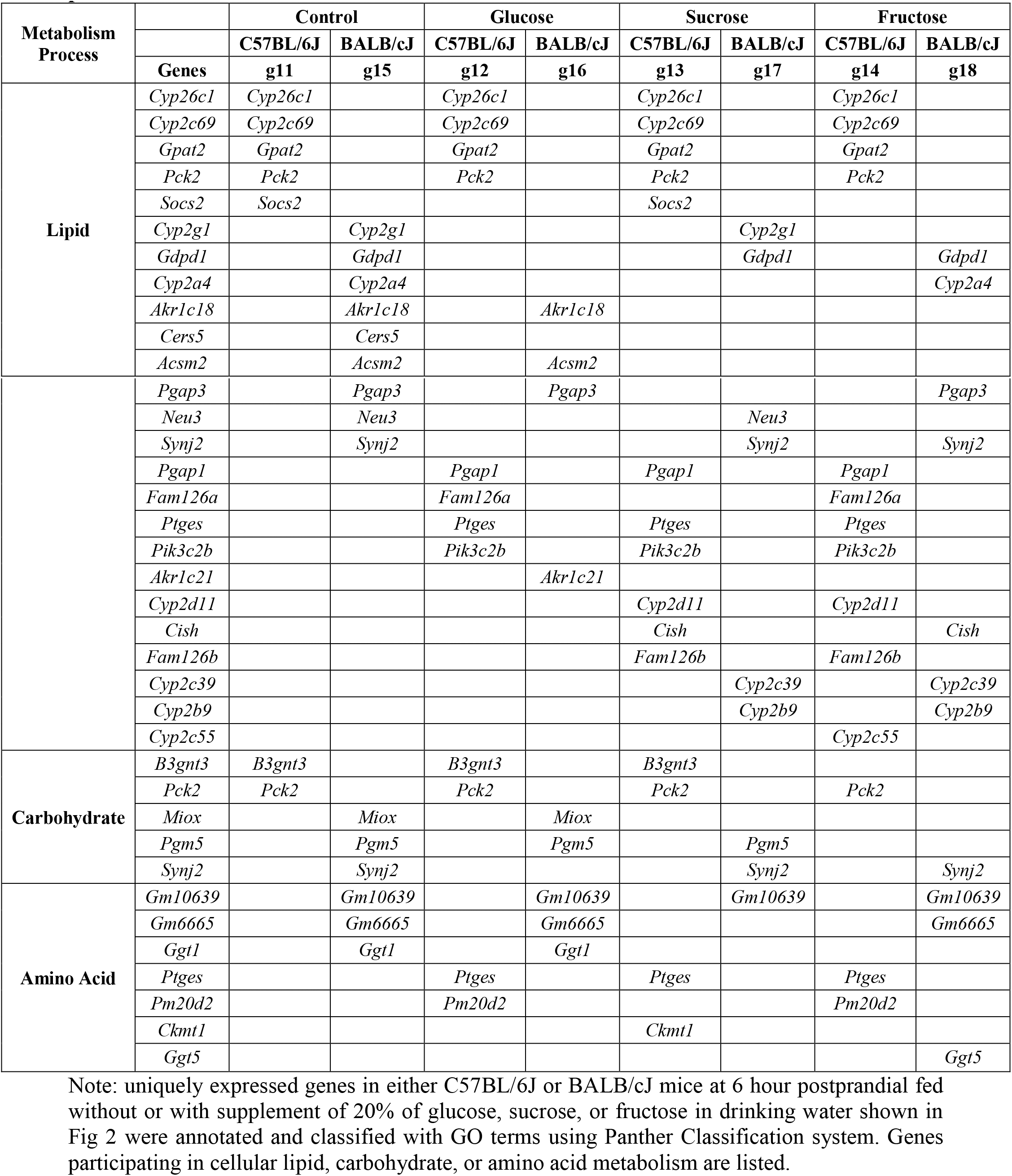
Genes uniquely expressed in two strains of mice participating in three metabolic processes.

As shown in Table 2, in both strains of mice and under all 4 conditions, there are more genes participating in lipid metabolism than in carbohydrate or amino acid metabolism. For lipid metabolism there are more genes uniquely expressed in BALB/cJ mice than in C57BL/6J mice when there was no carbohydrate supplement. However, supplement with any of those three carbohydrates resulted in more genes uniquely expressed in C57BL/6J mice than in BALB/cJ mice, suggesting that C57BL/6J mice are more sensitive to sugar intake than BALB/cJ mice. All three carbohydrates induced additional genes such as *Pgap1, Ptges, and Pik3c2b* in C57BL/6J mice, while diminished *Cers5* in BALB/cJ mice. As for the carbohydrate and amino acid metabolisms, supplements with those three carbohydrates caused some changes in gene expressions in both strains of mice.

To quantify the differential expression levels of genes between those two strains, we used DESeq2 R package to compare the expression level of each gene in BALB/cJ mice versus C57BL/6J mice and plotted −log10(p_adj_) versus log2(fold change) at 6 hour following feeding without or with different carbohydrate (Fig 3A-D). Genes with p_adj_ < 0.05 (-log10 (p_adj_) > 1.3) were viewed as significantly differentially expressed. Genes significantly highly expressed in C57BL/6J are in green and genes significantly highly expressed in BALB/cJ are in red. Genes with expression levels that were not significantly different between these two strains are in blue. Fig 3E summarizes the numbers of total significantly differentially expressed genes between two strains. In both strains, the numbers of differentially expressed genes are higher in control groups than in groups with any of those three carbohydrate supplements, suggesting that supplement with any of those carbohydrates diminished the differences in gene expression levels between those two strains of mice. Of those three carbohydrates, the numbers of differentially expressed genes and the degree of differences with fructose supplement are higher than those with either glucose or sucrose supplement.

**Fig 3.**
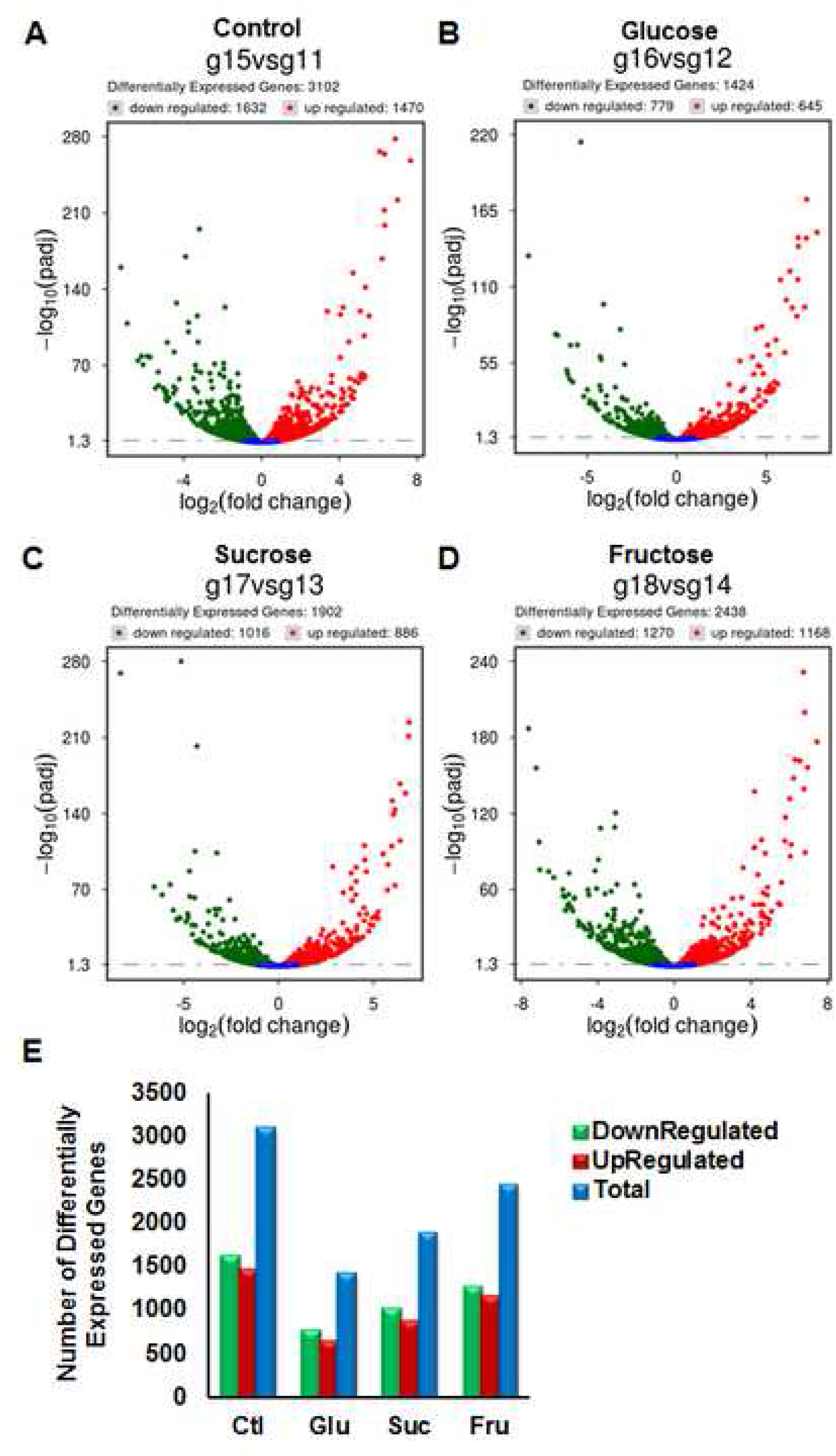
Quantification of differentially expressed genes in BALB/cJ mice versus C57BL/6J mice. (A-D) Volcano plots of −log10(p_adj_) versus log2(fold change) of genes expressed in BALB/cJ mice over those in C57BL/6J mice at 6 hour following feeding without or with different carbohydrates. p_adj_ and log2(fold change) were calculated using R package DESeq2. Significantly highly expressed genes in C57BL/6J mice are in green. Significantly highly expressed genes in BALB/cJ mice are in red. Genes with expression levels not significantly different between two stains are in blue. (E) Bar graph of total numbers of genes with significantly different expression levels between two strains.

We uploaded differentially expressed genes in green or red with threshold of |log2 (fold change)| ≥ 1 onto Panther Classification program to find out the biological functions of those genes by classification. Table 3 shows differentially expressed genes in two strains of mice participating in three metabolic processes. There are more genes in Table 3 than Table 2 and the majority of genes in lipid metabolic processes are different between these two tables, because large numbers of genes in Table 3 were from the shared populations rather than non-shared populations (Fig 2). Similar to Table 2, though, there are more genes involved in lipid metabolism than carbohydrate or amino acid metabolism. Sucrose and fructose induced more genes in C57BL/6J than BALB/cJ mice, again, suggesting that C57BL/6J mice are more sensitive to those carbohydrates than BALB/cJ mice.

**Table 3.**
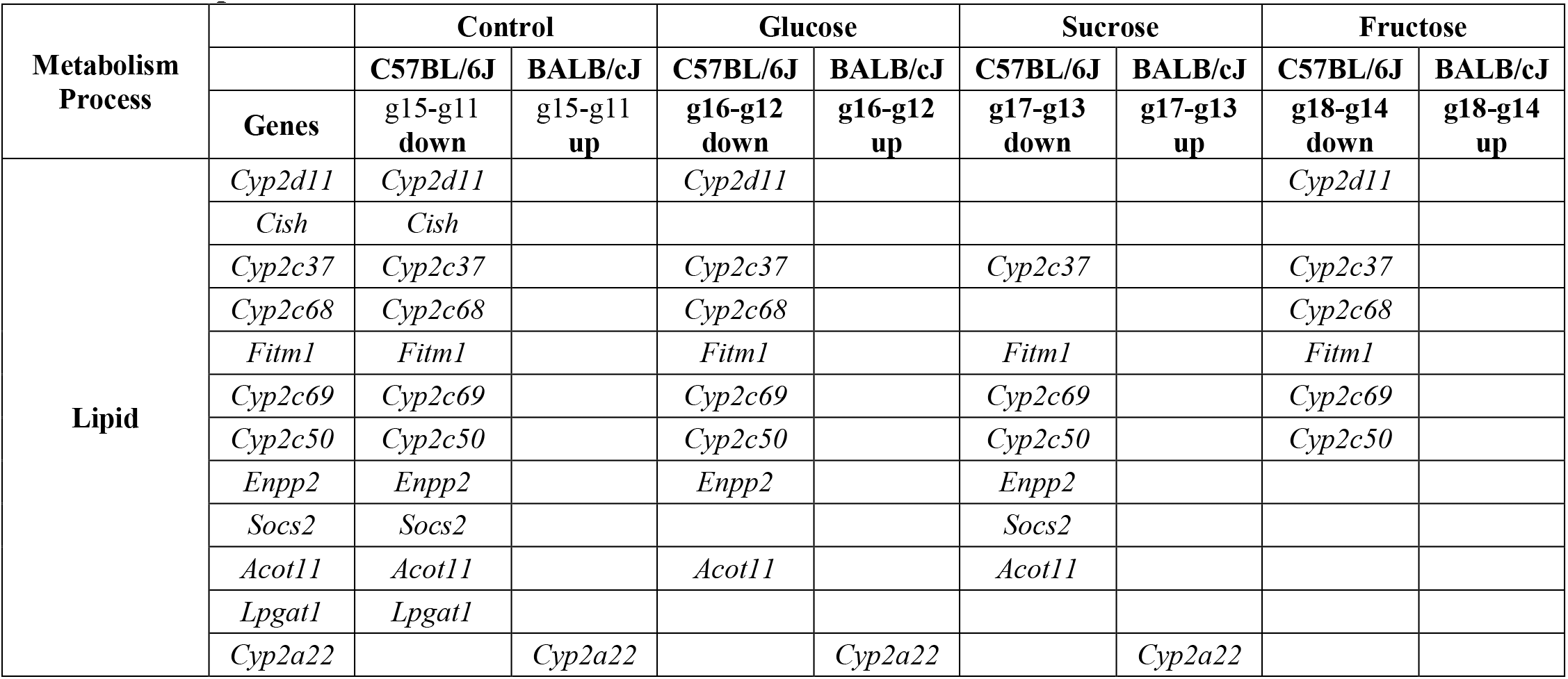
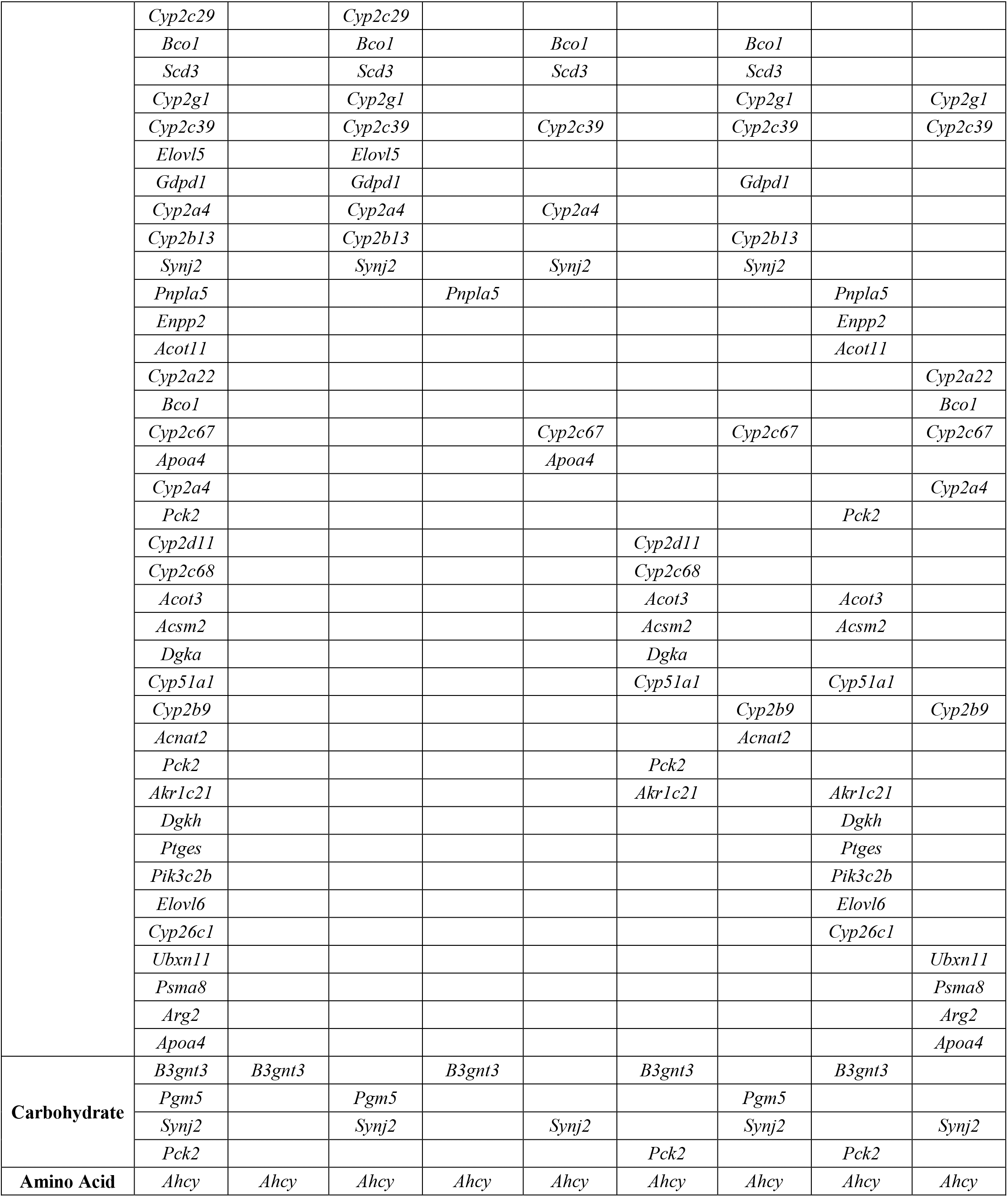
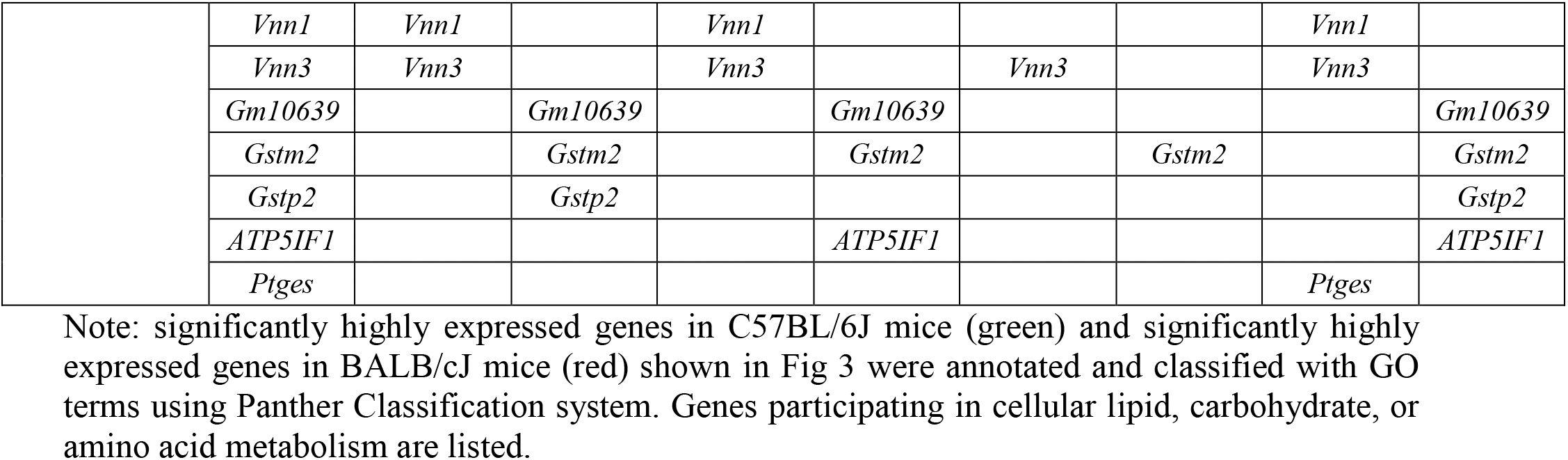
Differentially expressed genes in two stains of mice participating in three metabolic processes.

### Impact of dietary carbohydrate supplements on genome wide gene expression in two strains of mice

One evident result from the data shown thus far is that supplements with all three carbohydrates (glucose, sucrose and fructose) affected liver gene expressions and the effects were different among these three carbohydrates. To better understand the basis for these differences, we first determined whether carbohydrate supplemented water alter caloric intake. For C57Bl/6J mice, supplement with any of those three carbohydrates did not change drink intake (S2A Fig). However, compared to control, supplement with these carbohydrates individually resulted in extra 2.2 – 2.6 kcal intake, while there was no significant difference between glucose, sucrose or fructose (S2C Fig). In parallel, food intake was significantly reduced in mice supplemented with any of these three carbohydrates compared to the mice given plain water (S2B Fig). Correspondingly, the caloric intake from solid food was reduced in carbohydrate supplemented mice compared to mice given plain water, but again there was no significant difference between the three carbohydrate supplements (S2D Fig). Combining the total caloric intake (solid food and drinking), there was no significant difference between control and any of the carbohydrate water supplemented mice (S2E Fig). In contrast, BALB/cJ mice supplemented with either sucrose or fructose consumed less drink compared to mice given plain water, while glucose did affect the drink consumption (S2A Fig). These resulted in fewer calorie intake from drinking sucrose or fructose supplement water compared to glucose (S2A Fig), albeit all three carbohydrates increased calorie intake (S2C Fig). Despite this difference, the BALB/cJ mice all consumed similar amounts of calories from solid food independent of the additional calories present in drinking water (S2D Fig). Thus, the total caloric intake was greater for the mice ingesting glucose and fructose supplemented water compared to plain water, but did not reach statistical significance for the sucrose supplemented water group (S2E Fig). Although these differences are statistically significant, they are relatively small and both strains ingested between 7-8 kilocalories during the 3 h feeding period.

To visualize the overall gene expression changes we generated Venn diagrams showing shared and non-shared gene populations for the four groups, control (ctl), glucose (glu), sucrose (suc) and fructose (fru) groups at various time points postprandial in two strains of mice (Fig 4). At each time point, the numbers of genes shared by all 4 groups are in the range of 10,175 to 10,471. The numbers of genes in only one group that are not shared by other groups are in the range of 41 to 198. The numbers of genes shared by three carbohydrate groups are in the range of 34 to 182. Specific numbers of gene in 10 sets are listed in Fig 4G.

**Fig 4.**
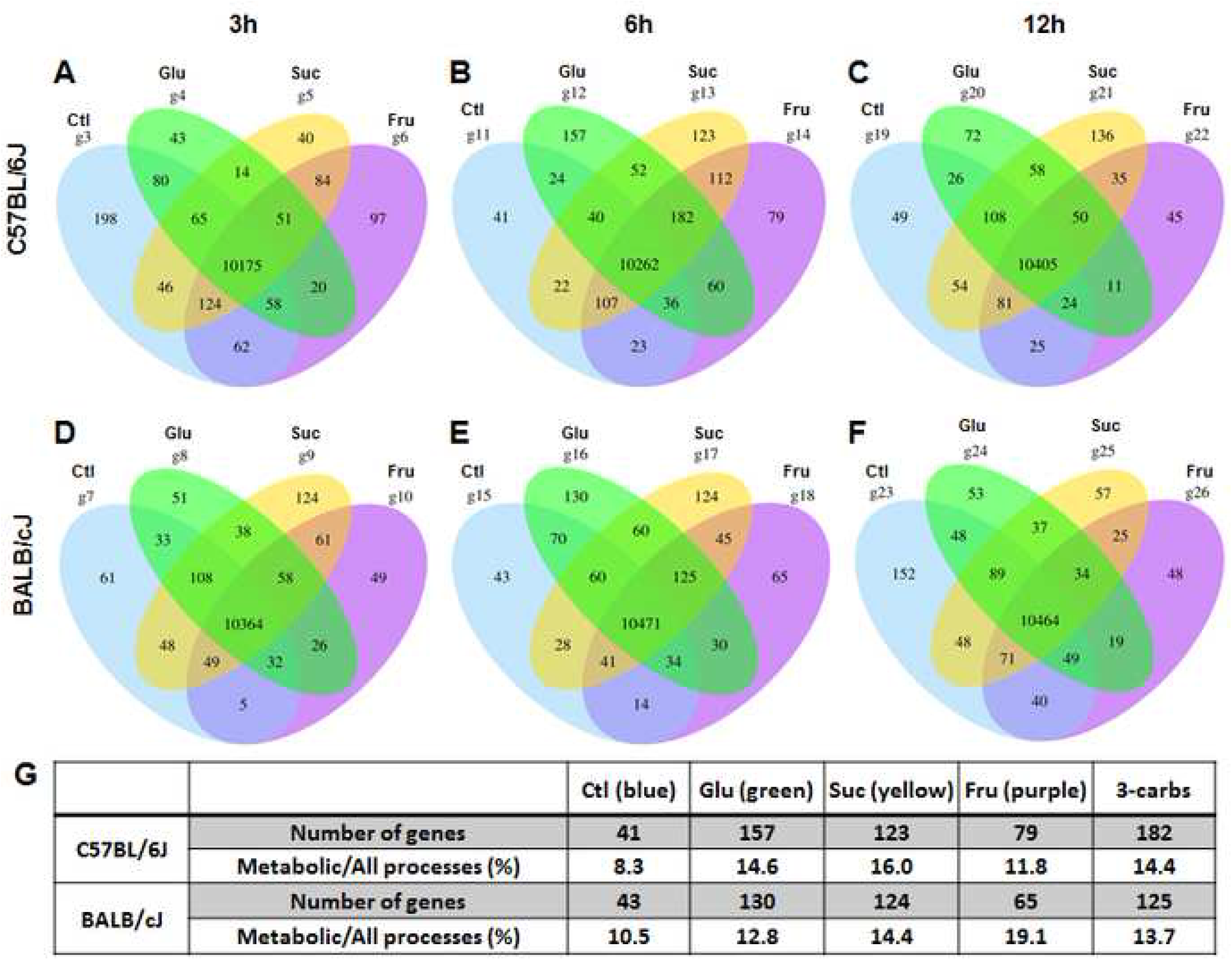
Gene distributions and populations in two strains of mice at various time points following feeding without or with carbohydrate supplements. Blue, control (Ctl, without carbohydrate supplement); green, yellow and purple, with supplement of 20% glucose, sucrose and fructose in drinking water, respectively. Genes with FPKM ≥ 1 were considered expressed. Genes with FPKM < 1 were considered absent. (G) Numbers of genes uniquely expressed groups fed without or with individual carbohydrate supplements and genes shared by three separate groups fed with individual supplement of glucose, sucrose or fructose, and percentages of genes participating in metabolic processes to all genes in corresponding populations.

We then uploaded these 10 sets of genes unto Panther Classification program and classified those genes according to their functions in biological processes. The percentages of genes involved in metabolic process to all biological processes are listed in Fig 4G and they are different among all sets. Supplement with each of these three carbohydrates increased the percentage of genes participating in metabolic processes in both strains by 20 – 100%. While sucrose caused the greatest increase in C57BL/6J mice, fructose caused the greatest increase in BALB/cJ mice. Specific genes participating in metabolic processes are listed in Table 4. The total number of genes participating in metabolic processes in all sets is 153. Of these 153 genes, 15 genes (10% of total) are shared by two strains of mice under the same or different conditions, indicating that the majority of genes are not shared by those strains. More importantly, the changes caused by any of those carbohydrates are different between two stains of mice. All three carbohydrates induced additional genes in both strains, and more genes were induced in C57BL/6J than in BALB/cJ mice, again, suggesting that C57BL/6J mice are more sensitive to the three carbohydrates than BALB/cJ mice. In C57BL/6J mice, glucose changed the highest number of genes compared to sucrose or fructose. In BABL/cJ mice, the numbers of changed genes caused by three individual carbohydrates are similar. The genes in the populations shared by three groups supplemented with three individual carbohydrates are the genes sensitive to any of three carbohydrates. These genes are different between two stains of mice, as only 1 (*Spidr*) out of more than 20 genes shared by both strains, demonstrating that the genes induced by all three individual carbohydrates are different between two strains of mice.

**Table 4.**
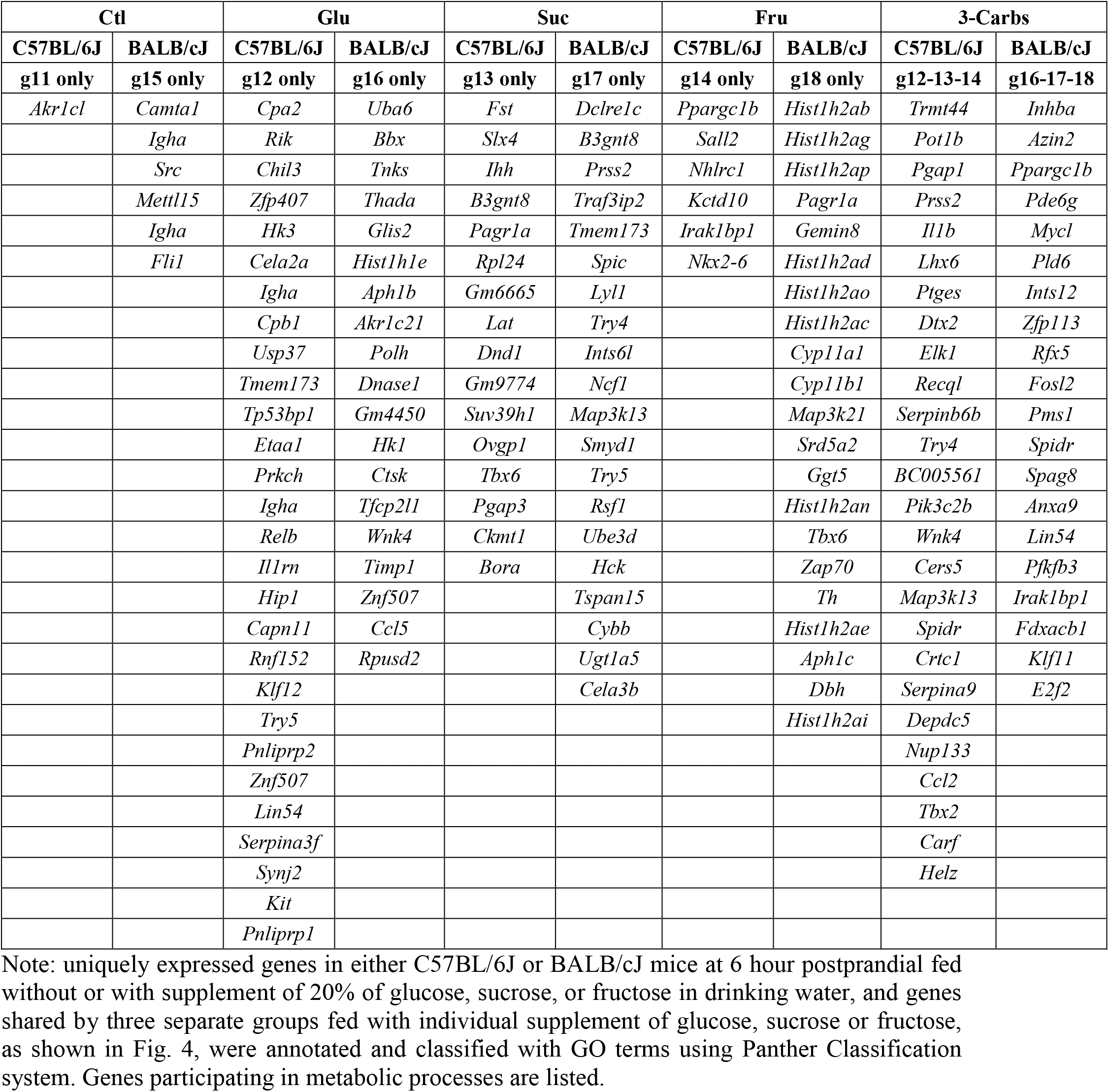
Uniquely expressed genes in different populations in two strains of mice participating in metabolic process.

To quantify those three individual carbohydrates caused changes in gene expression levels, we plotted −log10 (p_adj_) versus log2 (fold change) at 3, 6 and 12 hour following feeding without or with different carbohydrate (Figs 5–7). All three carbohydrates induced some genes and suppressed others in both strains. Again there were more genes affected by these three carbohydrates in C57BL/6J than in BALB/cJ mice. The greatest effects of glucose occurred at 12 hour postprandial in C57BL/6J mice and at 6 hour postprandial in BABL/cJ mice (Fig 5). With sucrose supplement, the highest number of changed genes occurred at 3 hour postprandial in both stains. Those numbers reduced rapidly in C57BL/6J mice than in BALB/cJ mice. In the case of fructose, the highest number of changed genes occurred at 3 hour postprandial in both strains and reduced thereafter. The reduction in C57BL/6J mice was not as rapid as with sucrose. Overall, fructose caused the greatest number of genes changed among the three carbohydrates.

**Fig 5.**
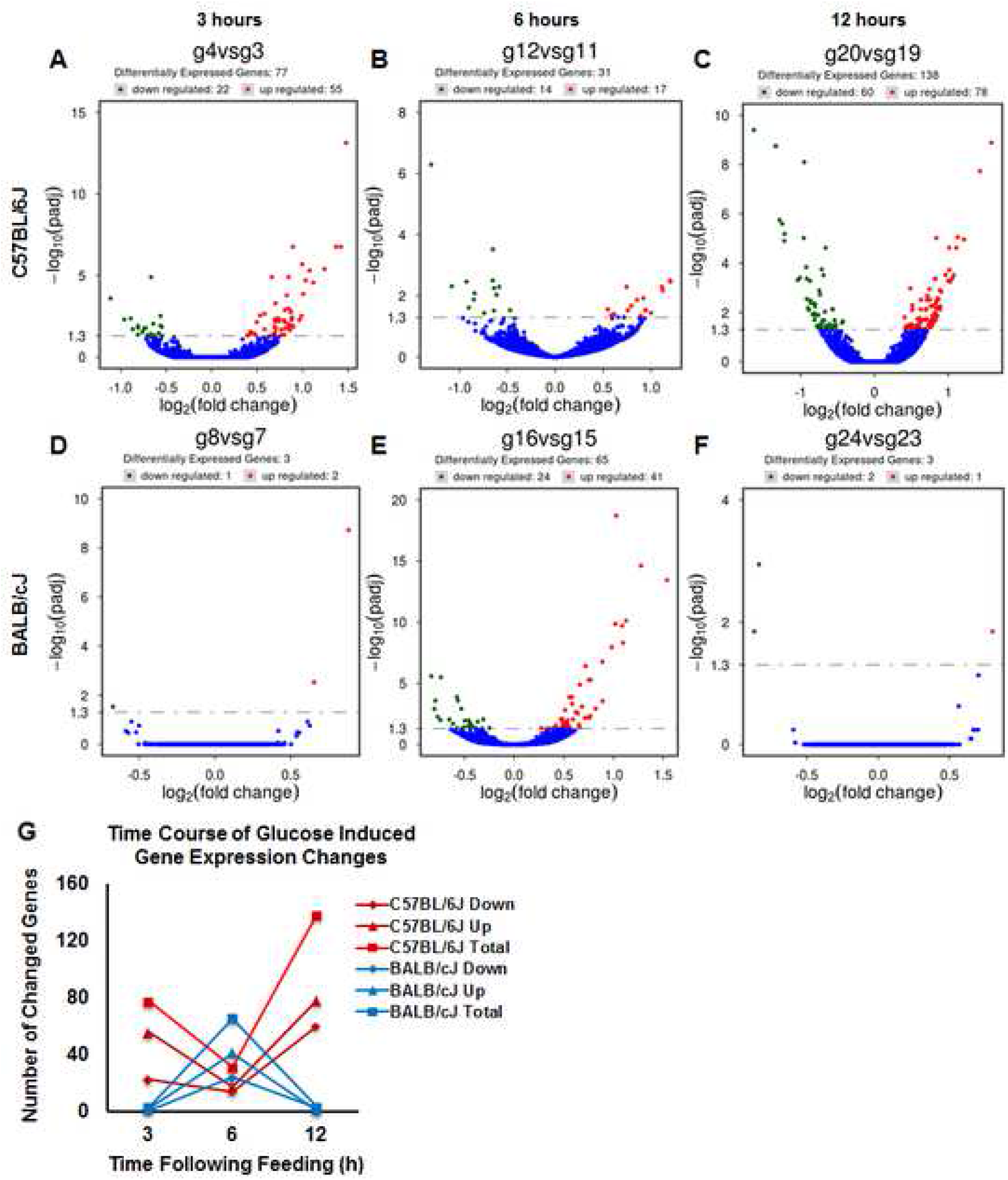
Glucose induced differentially expressed genes during the time course of 12 hours in two strains of mice. (A-F) Volcano plots of −log10(p_adj_) versus log2(fold change) at various time points following feeding. p_adj_ and log2(fold change) of all genes in C57BL/6J (A-C) or BALB/cJ (D-F) mice at 3 (A, D), 6 (B, E), or 12 (C, F) -hour time point following feeding supplemented with 20% glucose versus corresponding genes in controls (Ctls) (without any additional carbohydrate supplement) were calculated using R package DESeq2. Glucose significantly suppressed genes are in green. Glucose significantly induced genes are in red, genes not significantly changed by glucose are in blue. (G) Graph of number of glucose significantly changed genes.

**Fig 6.**
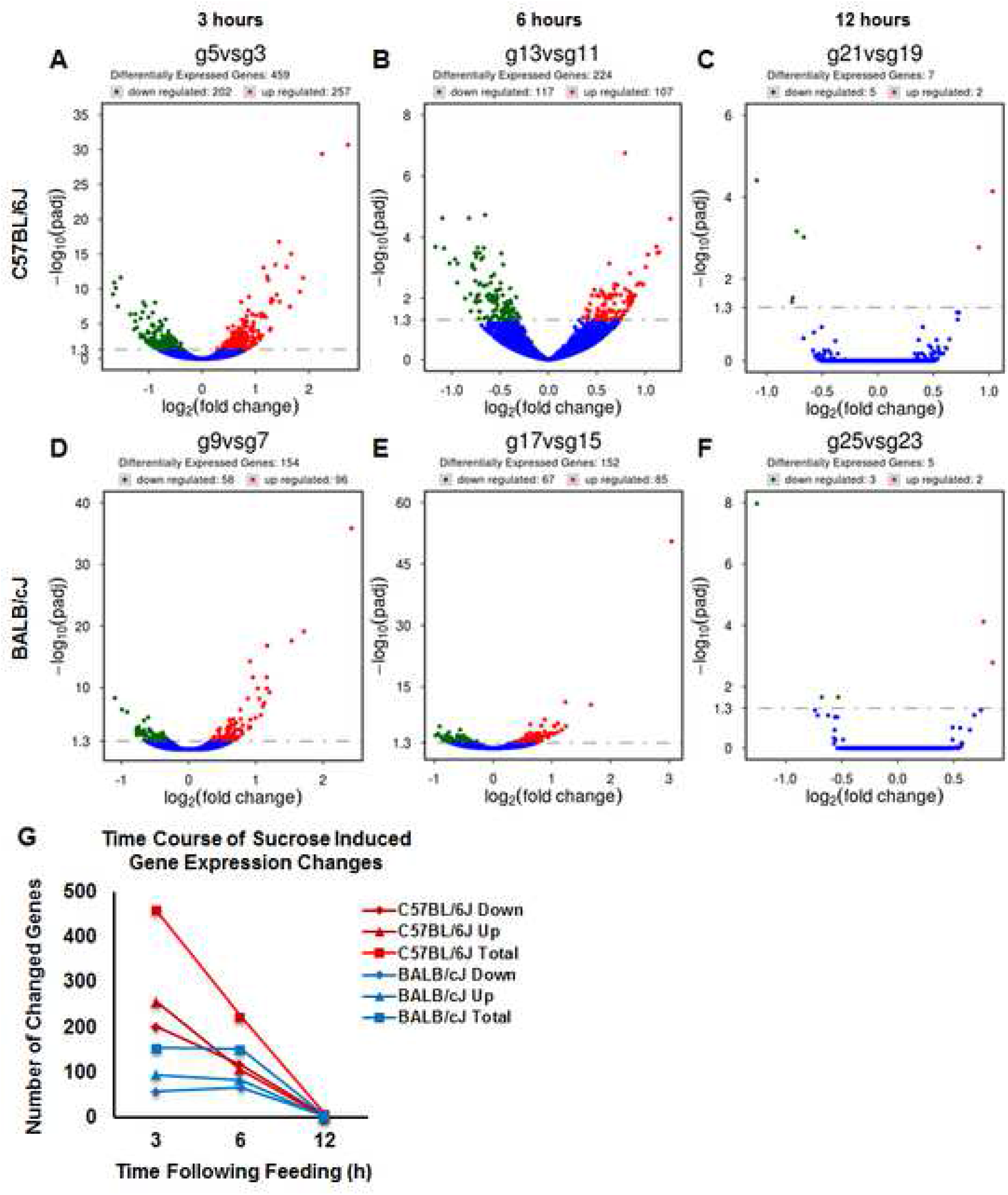
Sucrose induced differentially expressed genes during the time course of 12 hours in two strains of mice. (A-F) Volcano plots of −log10(p_adj_) versus log2(fold change) at various time points following feeding. p_adj_ and log2(fold change) of all genes in C57BL/6J (A-C) or BALB/cJ (D-F) mice at 3 (A, D), 6 (B, E), or 12 (C, F) -hour time point following feeding supplemented with 20% sucrose (B, E) versus corresponding genes in controls (Ctls) (without any additional carbohydrate supplement) were calculated using R package DESeq2. Sucrose significantly suppressed genes are in green. Sucrose significantly induced genes are in red, genes not significantly changed by glucose are in blue. (G) Graph of number of sucrose significantly changed genes.

**Fig 7.**
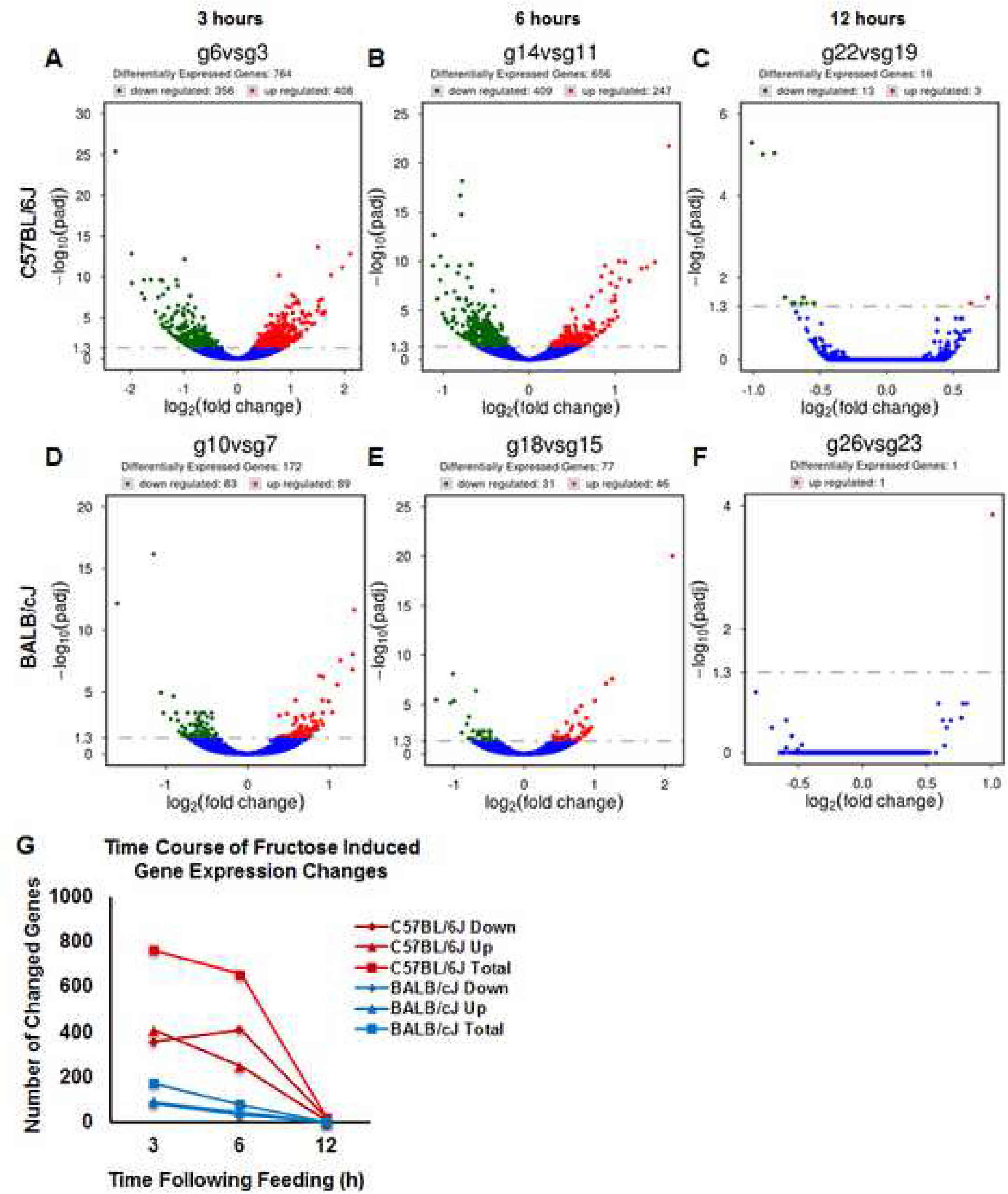
Fructose induced differentially expressed genes during the time course of 12 hours in two strains of mice. (A-F) Volcano plots of −log10(p_adj_) versus log2(fold change) at various time points following feeding. p_adj_ and log2(fold change) of all genes in C57BL/6J (A-C) or BALB/cJ (D-F) mice at 3 (A, D), 6 (B, E), or 12 (C, F) -hour time point following feeding supplemented with 20% fructose versus corresponding genes in controls (Ctls) (without any additional carbohydrate supplement) were calculated using R package DESeq2. Fructose significantly suppressed genes are in green. Fructose significantly induced genes are in red, genes not significantly changed by fructose are in blue. (G) Graph of number of glucose significantly changed genes.

To further investigate the causal relationships and to predict the signal pathways involved, we performed IPA of genes significantly changed by fructose. With fructose supplement and at 3 hour postprandial, 764 genes were significantly changed (356 suppressed and 408 induced) in C57BL/6J mice, while 172 genes were significantly changed (83 suppressed and 89 induced) in BALB/cJ mice. We uploaded those 764 genes of C57BL/6J mice and 172 genes of BALB/cJ mice separately in the IPA program and obtained two major networks for C57BL/6J mice (Fig 8A, B) and 1 network for BALB/cJ mice (Fig 8C). All of these networks are focused on lipid metabolisms, indicating that fructose changed genes mainly involved in lipid metabolisms in both strains. The fructose induced networks and signaling pathways are more complex in C57BL/6J than in BALB/cJ mice, as the analyses resulted in two distanced networks for C57BL/6J mice and only 1 network for BALB/cJ mice. The main players in network A are PPARα and PPARγ, both of which are important transcription factors regulating lipid metabolism, particularly lipid storage and synthesis [16]. Network B shows that several major players in lipid synthesis, including *Fasn, Acaca*, and their transcriptional factor *SREBP1* were upregulated by fructose in C57BL/6J mice. Network shown in C is similar to that in B, although there are more genes and signaling pathways upregulated in network B than C.

**Fig 8.**
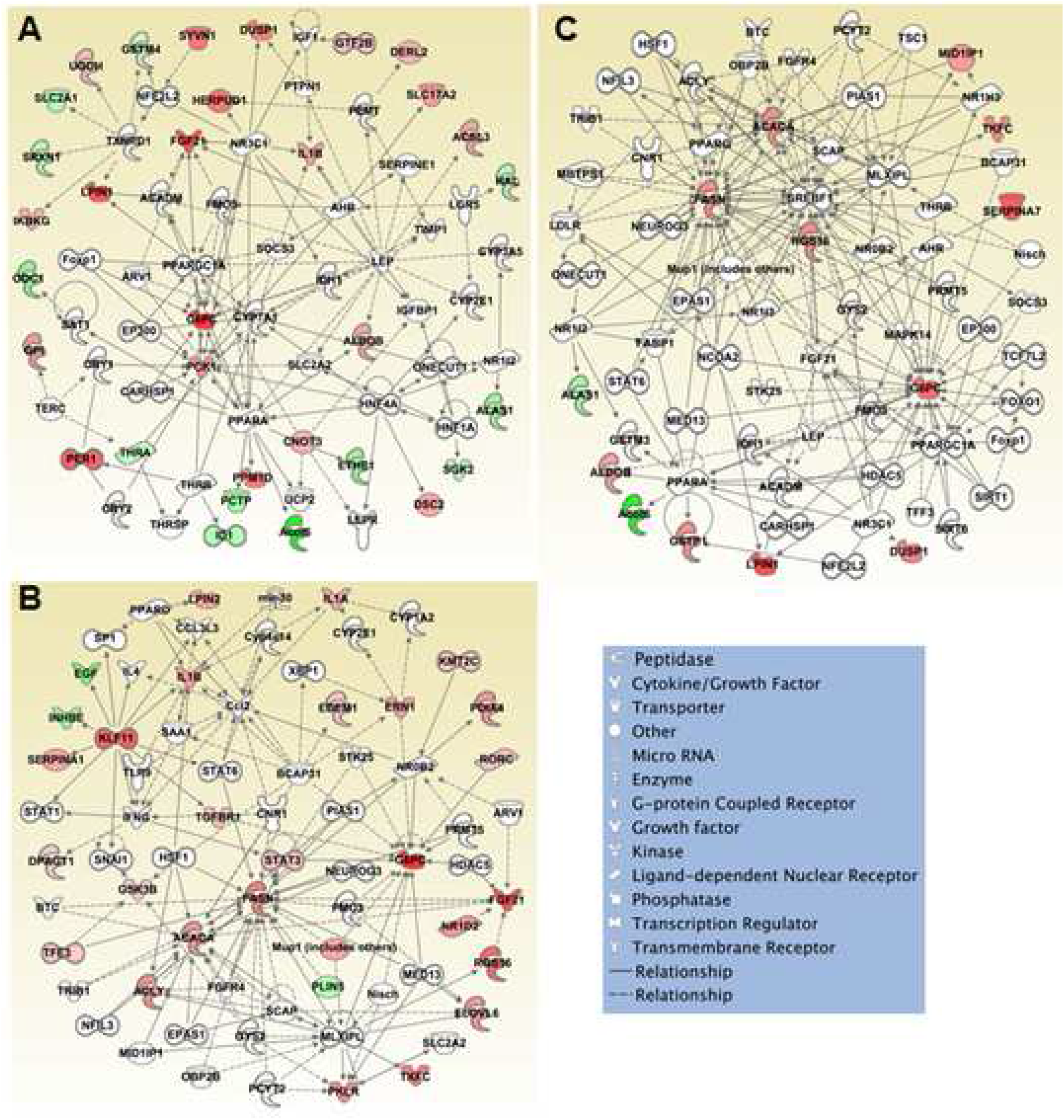
Intuitive Pathway Analysis (IPA) of networks of hepatic genes significantly changed by fructose and predicted signaling pathways in C57BL/6J (A, B) and BALB/cJ (C) mice. Solid lines are direct linked relationships and dotted lines are indirectly linked relationships. Green, down-regulated; red, up-regulated.

## Discussion

The liver plays an essential role in the control of glucose and lipid homeostasis. In the fasted state, the liver undergoes glycogenolysis and gluconeogenesis to release glucose in the systemic circulation for the maintenance of euglycemia [17, 18]. In contrast, the liver suppresses the *de novo* synthesis of fatty acids and the release of triglycerides as very low-density lipoprotein particles [18, 19]. These physiological responses are reverse in the fed state in which gluconeogenesis is inhibited and glycogen synthesis is increased, generating a net suppression of hepatic glucose output [18]. In parallel, dietary fatty acids and conversion of carbohydrates to fatty acids, *via de novo* lipogenesis are esterified into triglycerides for subsequent secretion as very low-density lipoprotein particles. The nutrient-dependent regulatory control of these pathways in the liver occurs at multiple levels including rapid and dynamic changes in gene expression [12, 20, 21].

Recently, we reported robust RNA-seq data sets for the liver of C57BL/6J and BALB/cJ mice in the fasted state and the time-dependent changes that occurred following feeding with a standard low-fat mouse chow [12]. However, consumption of food in western diets usually includes sweetened beverages. Prior to the 1950’s beverages were typically sweetened with sucrose (a disaccharide of glucose and fructose) but since then increasingly this was replaced with high fructose corn syrup (HFCS). These sugars have different routes of metabolism and in particular, fructose has been linked to obesity and the development of fatty liver disease [22, 23]. To provide a resource and a baseline for understanding the relative functional role of glucose, sucrose and fructose on liver gene expression, we analyzed liver mRNA gene expression under defined fasting and carbohydrate supplemented feeding states in two strains of mice.

Consistent with our previous data sets, these data demonstrate that the liver expresses over 10,000 mRNAs in both C57BL/6J and BALB/cJ mice, but that each has approximately 300-600 genes (depending upon dietary condition) that are uniquely expressed in one strain (Fig 2). Moreover, with the shared genes, the expression levels are different in one strain from the other (Fig 3). These differences led to greater inter-strain distances in gene expressions than the distances among intra-strain with different carbohydrates (Fig 1). Nevertheless, within each strain, supplements with different sugars caused significantly different gene expression patterns (Figs 4–7).

In this study, glucose, sucrose or fructose was individually given to mice via drinking water for only 3 hours when mice were switched from fasted to fed states on the experiment day. Given that the duration of high carbohydrate treatments was such a short period of time, the carbohydrate caused changes in gene expressions were impressively significant. Sucrose, the mostly consumed table sugar, is a disaccharide and its metabolism starts with hydrolysis to glucose and fructose. One would assume that the effects of sucrose are the combination of the effects of glucose and fructose. Interestingly, our in-depth analyses revealed that some genes were only expressed in sucrose treated mice and those sucrose induced unique genes were different between C57BL/6J and BALB/cJ mice (Fig 4, Tables 2, 4, S3). While whether sucrose could directly cause metabolic disorders is still debatable, several studies have shown that high sucrose intake increased triglyceride, cholesterol levels in human [2] and in animal models [24–26]. Here we found that several genes participating in lipid metabolism were altered (Tables 2–4, S3), suggesting that those altered genes could be responsible for sucrose caused hyperlipidemia. The alterations in C57BL/6J were different from those in BALB/cJ mice, which could partially explain why different strains or individuals have different responses to high sucrose intake. Glucose also uniquely induced genes that were not shared by either sucrose or fructose, which could signal unique glucose induced metabolic processes.

Compared to sucrose or glucose, fructose caused the greatest changes in gene expressions, and those changes were more profound in C57BL/6J mice than in BALB/cJ mice (Figs 5–7). In C57BL/6J mice fructose significantly increased expressions of genes playing important roles in fatty acid and lipid synthesis including elongation of very long chain fatty acids protein 6 (*Elovl6*) and ATP-citrate synthase (*Acly*), which could serve as molecular mechanisms of how fructose induced lipid accumulation in liver of nutrient sensitive animal models [27].

Besides these gene populations uniquely induced by each individual carbohydrates (Fig 4), there are genes that were shared by three groups treated with three individual carbohydrates (Fig 4, Tables 2, 4, S3), which could be the common genes mediating the signaling pathways to metabolic processes and physiological manifestations of the effects of all those three carbohydrates. Meanwhile, there are genes that were not affected by any of these three carbohydrates (Fig 4).

One might question whether carbohydrate supplement caused gene expression changes were indirect responses to changes in total energy intake and whether the unique changes caused by different carbohydrates were due to different energy intake as results of differernt carbohydrate supplements, rather than different types of carbohydrate. S2 Fig shows that there was no significant difference in total energy intake among control group and the groups supplemented with any of those three carbohydrates, especially fructose, in C57BL/6 J mice. In BALB/cJ mice, however, supplements with both glucose and fructose resulted in higher total energy intake compared to control. These results suggest that much greater gene expression changes in C57BL/6J mice, as compared to BALB/cJ mice, were not caused by total energy intake. Nutrient sensitive C57BL/6J mice were able to adjust their food intake when they were given extra energy via drinking. Fructose induced the greatest gene expression changes in C57BL/6J mice (Fig 3, Figs 5–8). Yet, total energy intake with fructose supplement was almost the same as that without any carbohydrate supplement, suggesting that gene expression changes cannot be attributed to total energy intake.

Overall, both strains displayed significant differences in their responses to the three carbohydrates in terms of the magnitude and rate of gene changes as well as in the engagement of signaling networks. For example, although fructose altered the expression of the greatest number of mRNAs with a similar rate in two strains of mice, the extent of changes was much greater in the C57BL/6J mice than in BABL/cJ mice (Fig 7). In contrast, the maximal change induced by glucose occurred at 12 h postprandial in C57BL/6J mice but at 6 h postprandial in BABL/cJ mice (Fig 5). While the sucrose stimulated time frame of mRNA induction was similar in the two strains, the rate of return to the fasted state expression levels was faster in C57BL/6J than in BALB/cJ mice (Fig 6).

Another interesting distinction between C57BL/6J than BALB/cJ mice is the network pathways activated by fructose. In C57BL/6J mice, two separate networks were identified that regulate different aspects of lipid metabolism, one centered around the peroxisome proliferator-activated receptors, PPARα and PPARγ, important for lipid storage and utilization, and the second network around lipid synthesis, SREBP and MLXIPL. In contrast, fructose primarily induced the network driving lipid synthesis in BALB/cJ mouse livers. The differences in network regulation may help to explain the relative resistance of BALB/cJ mice to diet induced insulin resistance and obesity compared to C57BL/6J mice.

## Conclusions

In summary, the data presented in this manuscript provide a robust data resource for the acute effects of three common dietary sugars on the liver transcriptomes of C57BL/6J and BALB/cJ mice. These data demonstrate that each of these sugars activates qualitatively and quantitatively different patterns of gene expression upon feeding and during the subsequent return to the fasted state. In addition, this unique data resource will provide a framework to which other studies can now be compared.

## Supporting information

**S1 Table. Data quality summary.**

**S2 Table. Correlation coefficients between one and all other samples in each of all 26**

**groups.**

**S3 Table Genes uniquely expressed in two strains of mice under different conditions participating in metabolic processes.**

**S1 Fig. Correlation of gene expressions.** Correlation matrix of all 128 samples in 26 groups.

**S2 Fig. Drink, food and calorie intake.** Drink intake (A), food intake (B), and calorie intake via drinking (C) or food (D), or total calorie intake (C) of mice during the first 3 hours when 20% of glucose, sucrose, or fructose was separately supplemented in drinking water, n = 7-9 per group. Value are mean ± S.D. p values were obtained by two tail student t-test. *p < 0.05, **p < 0.01, or ***p < 0.001 was considered significant. Unit calories: glucose, 3.8 kcal/g; sucrose, 3.9 kcal/g; fructose 3.6 kcal/g; food, 4.11 kcal/g.

## Notes

### Competing Interest Statement

The authors have declared no competing interest.

