## Supplementary Materials for "The regulation of liver gene expression by carbohydrates is mouse strain specific"

**S1 Table. Data quality summary**

| Sample name | Raw reads | Clean reads | Raw bases | Clean bases | Error rate (%) | Q20 (% ) | Q30 (%) | GC content (%) |
|---|---|---|---|---|---|---|---|---|
| JC001 | 61941814 | 59220206 | 9.3G | 8.9G | 0.02 | 97.94 | 94.40 | 49.84 |
| JC002 | 74601006 | 71400296 | 11.2G | 10.7G | 0.03 | 97.74 | 93.98 | 50.52 |
| JC003 | 71476110 | 68217992 | 10.7G | 10.2G | 0.02 | 97.94 | 94.37 | 49.29 |
| JC004 | 82036442 | 78410590 | 12.3G | 11.8G | 0.03 | 97.91 | 94.30 | 50.06 |
| JC005 | 70923404 | 67843458 | 10.6G | 10.2G | 0.03 | 97.81 | 94.07 | 49.37 |
| JC006 | 70903168 | 68529728 | 10.6G | 10.3G | 0.03 | 97.92 | 94.29 | 50.08 |
| JC007 | 73183310 | 70726372 | 11G | 10.6G | 0.03 | 97.86 | 94.17 | 50.38 |
| JC008 | 74509502 | 72222124 | 11.2G | 10.8G | 0.03 | 97.91 | 94.31 | 50.57 |
| JC009 | 68248824 | 65894208 | 10.2G | 9.9G | 0.03 | 97.90 | 94.32 | 50.30 |
| JC010 | 70395082 | 66907562 | 10.6G | 10G | 0.02 | 97.94 | 94.39 | 50.58 |
| JC011 | 74292796 | 70527168 | 11.1G | 10.6G | 0.02 | 97.98 | 94.48 | 48.99 |
| JC012 | 82038278 | 78674894 | 12.3G | 11.8G | 0.03 | 97.68 | 93.89 | 49.20 |
| JC013 | 82831678 | 79690386 | 12.4 G | 12G | 0.03 | 97.67 | 93.79 | 49.82 |
| JC014 | 68620746 | 65699466 | 10.3G | 9.9G | 0.03 | 97.63 | 93.71 | 49.37 |
| JC015 | 70940914 | 67963478 | 10.6G | 10.2G | 0.03 | 96.49 | 91.31 | 49.48 |
| JC016 | 64490612 | 61611044 | 9.7G | 9.2G | 0.03 | 97.51 | 93.49 | 48.78 |
| JC017 | 64055914 | 61077736 | 9.6G | 9.2G | 0.03 | 97.40 | 93.32 | 49.48 |
| JC018 | 60387300 | 57303212 | 9.1G | 8.6G | 0.03 | 97.31 | 93.04 | 50.36 |
| JC019 | 63605456 | 59784152 | 9.5G | 9G | 0.03 | 97.57 | 93.56 | 49.27 |
| JC020 | 61592410 | 59473184 | 9.2G | 8.9G | 0.03 | 97.36 | 93.13 | 49.20 |
| JC021 | 65086574 | 62189648 | 9.8G | 9.3G | 0.03 | 97.50 | 93.39 | 48.53 |
| JC022 | 72180810 | 69560234 | 10.8G | 10.4G | 0.03 | 97.46 | 93.18 | 49.72 |
| JC023 | 72586792 | 69458842 | 10.9G | 10.4G | 0.03 | 97.38 | 93.18 | 49.94 |

| | | | | | | | | |
|---|---|---|---|---|---|---|---|---|
| JC024 | 71478072 | 67561766 | 10.7G | 10.1G | 0.03 | 97.42 | 93.20 | 48.50 |
| JC025 | 66327372 | 62534608 | 9.9G | 9.4G | 0.03 | 97.54 | 93.59 | 50.39 |
| JC026 | 62129594 | 58702410 | 9.3G | 8.8G | 0.03 | 97.34 | 93.10 | 50.09 |
| JC027a | 90470880 | 88733034 | 13.6G | 13.3G | 0.02 | 98.44 | 95.25 | 48.28 |
| JC028 | 65968054 | 63514270 | 9.9G | 9.5G | 0.03 | 97.28 | 92.97 | 49.19 |
| JC029a | 74317096 | 71841168 | 11.1G | 10.8G | 0.03 | 97.86 | 94.12 | 49.60 |
| JC030a | 72137620 | 70060922 | 10.8G | 10.5G | 0.02 | 98.43 | 95.27 | 48.18 |
| JC031 | 74335166 | 69189442 | 11.2G | 10.4 | 0.03 | 97.23 | 92.79 | 48.48 |
| JC032 | 64226726 | 61757970 | 9.6G | 9.3G | 0.03 | 97.27 | 92.89 | 48.60 |
| JC033 | 76058264 | 74 969638 | 11.4G | 11.2G | 0.03 | 97.52 | 93.49 | 49.80 |
| JC034a | 73158936 | 71311364 | 11G | 10.7G | 0.02 | 98.48 | 95.46 | 48.84 |
| JC035a | 84264858 | 82059344 | 12.6G | 12.3G | 0.02 | 98.48 | 95.44 | 49.01 |
| JC036a | 82329306 | 80467874 | 12.3G | 12.1G | 0.02 | 98.51 | 95.52 | 48.89 |
| JC037a | 81010026 | 78983026 | 12.2G | 11.8G | 0.02 | 98.61 | 95.73 | 47.89 |
| JC038a | 89904876 | 87558378 | 13.5G | 13.1G | 0.02 | 98.50 | 95.48 | 47.77 |
| JC039 | 64451310 | 60191242 | 9.7G | 9G | 0.03 | 97.41 | 93.22 | 48.43 |
| JC040a | 72763844 | 70773198 | 10.9G | 10.6G | 0.02 | 98.32 | 94.99 | 47.97 |
| JC041a | 71954564 | 69982214 | 10.8G | 10.5G | 0.02 | 98.31 | 94.99 | 48.69 |
| JC042 | 64160286 | 61621362 | 9.6G | 9.2G | 0.03 | 97.48 | 93.40 | 48.60 |
| JC043a | 72378194 | 70340878 | 10.9G | 10.6G | 0.02 | 98.53 | 95.57 | 48.60 |
| JC044a | 74229664 | 72480566 | 11.1G | 10.9G | 0.03 | 97.40 | 92.94 | 48.68 |
| JC04 5a | 75520374 | 73599082 | 11.3G | 11G | 0.02 | 98.38 | 95.14 | 48.56 |
| JC046a | 66211004 | 64497972 | 9.9G | 9.7G | 0.02 | 98.47 | 95.38 | 48.62 |
| JC047a | 61070476 | 59502474 | 9.2G | 8.9G | 0.03 | 97.48 | 93.10 | 48.53 |
| JC048a | 88272246 | 85927566 | 13.2G | 12.9G | 0.02 | 98.31 | 94 .97 | 48.89 |
| JC049a | 70762088 | 69366184 | 10.6G | 10.4G | 0.02 | 98.46 | 95.49 | 49.25 |

| | | | | | | | | |
|---|---|---|---|---|---|---|---|---|
| JC050a | 83491152 | 81954934 | 12.5G | 12.3G | 0.02 | 98.50 | 95.55 | 49.00 |
| JC051a | 89123394 | 87240712 | 13.4G | 13.1G | 0.02 | 98.40 | 95.28 | 48.79 |
| JC052a | 9514 8612 | 93539586 | 14 .3G | 14 G | 0.02 | 98.46 | 95.43 | 48.70 |
| JC053 | 68699352 | 66446424 | 10.3G | 10G | 0.03 | 97.43 | 93.28 | 49.86 |
| JC054 | 66129434 | 63591382 | 9.9G | 9.5G | 0.03 | 97.53 | 93.56 | 50.54 |
| JC055 | 62805872 | 59272320 | 9.4G | 8.9G | 0.03 | 97.68 | 93.79 | 49.64 |
| JC056 | 75994268 | 71759146 | 11.4G | 10.8G | 0.03 | 97.52 | 93.49 | 49.53 |
| JC057 | 84056214 | 78590424 | 12.6G | 11.8G | 0.03 | 97.39 | 93.11 | 49.71 |
| JC058 | 62048778 | 58906968 | 9.3G | 8.8G | 0.03 | 97.44 | 93.31 | 50.03 |
| JC059 | 63607244 | 6114 9714 | 9.5G | 9.2G | 0.03 | 97.57 | 93.58 | 50.70 |
| JC060 | 63875984 | 61298914 | 9.6G | 9.2G | 0.03 | 97.10 | 92.36 | 49.71 |
| JC061 | 67568662 | 66456572 | 10.1G | 10G | 0.03 | 97.62 | 93.68 | 49.54 |
| JC062 | 76418606 | 71657240 | 11.5G | 10.7G | 0.03 | 97.65 | 93.78 | 49.27 |
| JC063 | 66220132 | 62110184 | 9.9G | 9.3G | 0.03 | 97.70 | 93.82 | 48.73 |
| JC064 | 119765114 | 117873082 | 18G | 17.7G | 0.02 | 98.11 | 94.68 | 48.81 |
| JC065 | 62634468 | 60214772 | 9.4G | 9G | 0.03 | 97.57 | 93.58 | 49.39 |
| JC066 | 63292690 | 60828288 | 9.5G | 9.1G | 0.03 | 97.60 | 93.64 | 49.50 |
| JC067 | 73357042 | 70092958 | 11G | 10.5G | 0.03 | 97.45 | 93.26 | 50.24 |
| JC068 | 71057770 | 67155478 | 10.7G | 10.1G | 0.03 | 97.49 | 93.44 | 50.28 |
| JC069 | 65924960 | 63266488 | 9.9G | 9.5G | 0.03 | 97.27 | 92.99 | 50.20 |
| JC070 | 69186252 | 63361340 | 10.4G | 9.5G | 0.03 | 97.61 | 93.66 | 49.79 |
| JC071 | 71010434 | 65464402 | 10.7G | 9.8G | 0.03 | 97.65 | 93.84 | 49.02 |
| JC072 | 74 642290 | 69555562 | 11.2G | 10.4G | 0.03 | 97.41 | 93.25 | 49.15 |
| JC073 | 61990826 | 57720260 | 9.3G | 8.7G | 0.03 | 97.45 | 93.41 | 49.62 |
| JC074 | 62410630 | 58522698 | 9.4G | 8.8G | 0.03 | 97.19 | 92.62 | 49.64 |
| JC075 | 65615532 | 61247284 | 9.8G | 9.2G | 0.03 | 97.67 | 93.77 | 49.13 |

| | | | | | | | | |
|---|---|---|---|---|---|---|---|---|
| JC076 | 70255274 | 64801940 | 10.5G | 9.7G | 0.03 | 97.46 | 93.36 | 50.34 |
| JC077 | 63877248 | 59807962 | 9.6G | 9G | 0.03 | 97.64 | 93.73 | 49.32 |
| JC078 | 63763520 | 61336346 | 9.6G | 9.2G | 0.03 | 97.61 | 93.64 | 51.23 |
| JC079 | 67775956 | 63237652 | 10.2G | 9.5G | 0.03 | 97.65 | 93.80 | 48.35 |
| JC080 | 76779628 | 71477494 | 11.5G | 10.7G | 0.03 | 97.26 | 93.03 | 50.01 |
| JC081 | 68755174 | 65947312 | 10.3G | 9.9G | 0.03 | 97.36 | 93.20 | 48.79 |
| JC082 | 65364528 | 61883414 | 9.8G | 9.3G | 0.03 | 97.47 | 93.39 | 48.81 |
| JC083a | 81508078 | 80089968 | 12.2G | 12G | 0.02 | 98.36 | 95.16 | 48.01 |
| JC084 | 62167006 | 58708766 | 9.3G | 8.8G | 0.03 | 97.30 | 93.06 | 48.01 |
| JC085 | 80264368 | 75299354 | 12G | 11.3G | 0.03 | 97.49 | 93.46 | 49.03 |
| JC086 | 64 934810 | 60744098 | 9.7G | 9.1G | 0.03 | 97.60 | 93.70 | 49.94 |
| JC087 | 64 87504 8 | 59985638 | 9.7G | 9G | 0.03 | 97.34 | 93.12 | 49.64 |
| JC088 | 66402050 | 62120902 | 10G | 9.3G | 0.03 | 97.59 | 93.61 | 50.07 |
| JC089 | 66812642 | 61746026 | 10G | 9.3G | 0.03 | 97.56 | 93.55 | 49.19 |
| JC090 | 63758652 | 61291272 | 9.6G | 9.2G | 0.03 | 97.57 | 93.62 | 48.98 |
| JC091 | 60098226 | 57629872 | 9G | 8.6G | 0.03 | 97.44 | 93.32 | 49.18 |
| JC092 | 63643886 | 60789686 | 9.5G | 9.1G | 0.03 | 97.11 | 92.57 | 48.86 |
| JC093 | 75423398 | 72024594 | 11.3G | 10.8G | 0.03 | 97.51 | 93.48 | 50.13 |
| JC094 | 62341216 | 58909280 | 9.4 G | 8.8G | 0.03 | 97.55 | 93.58 | 50.03 |
| JC095 | 76951642 | 73669502 | 11.5G | 11.1G | 0.03 | 97.69 | 93.77 | 50.17 |
| JC096 | 80258132 | 75647750 | 12G | 11.3G | 0.03 | 97.64 | 93.68 | 50.08 |
| JC097 | 65353814 | 62912378 | 9.8G | 9.4G | 0.03 | 97.46 | 93.31 | 46.79 |
| JC098 | 85323676 | 80144100 | 12.8G | 12G | 0.03 | 97.27 | 92.88 | 47.26 |
| JC099a | 80908072 | 79263042 | 12.1G | 11.9G | 0.02 | 98.49 | 95.49 | 48.09 |
| JC100 | 89725706 | 84 944752 | 13.5G | 12.7G | 0.03 | 97.39 | 93.12 | 46.74 |
| JC101 | 97418068 | 92376148 | 14 .6G | 13.9G | 0.03 | 97.35 | 93.10 | 48.51 |

| | | | | | | | | |
|---|---|---|---|---|---|---|---|---|
| JC103 | 87762618 | 83578780 | 13.2G | 12.5G | 0.03 | 97.23 | 92.82 | 49.16 |
| JC105 | 60567020 | 58715932 | 9.1G | 8.8G | 0.03 | 97.50 | 93.34 | 48.04 |
| JC106a | 66200188 | 63926592 | 9.9G | 9.6G | 0.03 | 97.67 | 93.50 | 49.28 |
| JC107a | 81084268 | 79596516 | 12.2G | 11.9G | 0.02 | 98.32 | 95.06 | 48.26 |
| JC108 | 73286700 | 70956132 | 11G | 10.6G | 0.03 | 97.29 | 92.96 | 47.73 |
| JC109a | 79139370 | 77676038 | 11.9G | 11.7G | 0.02 | 98.46 | 95.40 | 48.25 |
| JC110a | 76197856 | 74735020 | 11.4 G | 11.2G | 0.02 | 98.37 | 95.17 | 48.12 |
| JC111a | 85927750 | 84204662 | 12.9G | 12.6G | 0.02 | 98.43 | 95.34 | 48.84 |
| JC112 | 73868060 | 71041166 | 11.1G | 10.7G | 0.03 | 97.56 | 93.57 | 47.85 |
| JC113a | 80586500 | 79093658 | 12.1G | 11.9G | 0.02 | 98.27 | 94.97 | 48.76 |
| JC114 a | 76393082 | 74823372 | 11.5G | 11.2G | 0.02 | 98.36 | 95.24 | 48.87 |
| JC115 | 73531874 | 71367684 | 11G | 10.7G | 0.03 | 97.52 | 93.45 | 47.73 |
| JC116 | 70904448 | 68533032 | 10.6G | 10.3G | 0.03 | 97.93 | 94.23 | 48.77 |
| JC117a | 71829012 | 69950950 | 10.8G | 10.5G | 0.03 | 97.43 | 93.01 | 48.18 |
| JC118a | 83369624 | 81662312 | 12.5G | 12.2G | 0.02 | 98.43 | 95.42 | 47.68 |
| JC119 | 69820582 | 67229700 | 10.5G | 10.1G | 0.03 | 97.59 | 93.60 | 48.62 |
| JC120 | 71706180 | 68865362 | 10.8G | 10.3G | 0.03 | 97.42 | 93.30 | 48.4 1 |
| JC121 | 80207504 | 77859696 | 12G | 11.7G | 0.03 | 97.18 | 92.75 | 48.38 |
| JC122 | 84957064 | 81463418 | 12.7G | 12.2G | 0.03 | 97.49 | 93.42 | 48.03 |
| JC123a | 86084560 | 84 431888 | 12.9G | 12.7G | 0.02 | 98.21 | 94.79 | 47.81 |
| JC124 a | 87555400 | 85748980 | 13.1G | 12.9G | 0.02 | 98.40 | 95.26 | 47.64 |
| JC125a | 71547064 | 69899918 | 10.7G | 10.5G | 0.02 | 98.20 | 94.82 | 48.09 |
| JC126 | 78962390 | 76253618 | 11.8G | 11.4G | 0.03 | 96.02 | 90.27 | 47.66 |
| JC127 | 75601048 | 73517516 | 11.3G | 11G | 0.03 | 97.53 | 93.49 | 48.32 |
| JC128 | 65840508 | 63538778 | 9.9G | 9.5G | 0.03 | 97.32 | 93.12 | 48.85 |
| JC129 | 62069016 | 60090426 | 9.3G | 9G | 0.03 | 97.19 | 92.77 | 48.4 0 |

| | | | | | | | | |
|---|---|---|---|---|---|---|---|---|
| JC130 | 61887248 | 59433304 | 9.3G | 8.9G | 0.03 | 97.19 | 92.77 | 48.36 |

(1) Sample name: sample ID.
(2) Raw reads: reads count from the raw data, four rows as a unit, with statistics of reads count for every sequencing.
(3) Clean reads: reads count filtered from raw data. Statistics method is similar to raw reads. All the following analyses are based on clean data.
(4) Raw bases: Base number of raw data, (number of raw reads) * (sequence length), converting unit to G.
(5) Clean bases: Base number of raw data after filtering, (number of clean reads) * (sequence length), converting unit to G.
(6) Error rate (%): base error rate of whole sequencing.
(7) Q20 (%): Phred values greater than 20 base number contain the percentage of total bases, (base number of Phred value > 20) / (total base number)*100.
(8) Q30 (%): Phred values greater than 30 base number contain the percentage of total bases, (base number of Phred value > 30) / (total base number)*100.
(9) GC content (%): The percentage of G&C base numbers of total bases (G&C base number) / (total base number)*100.

**S2 Table. Correlation coefficients between one and all other samples in each of all 26 groups.**

| Group | Samples | Correlation | Correlation Coefficient ($R^2$) |
|---|---|---|---|
| g1 | JC001 JC027a JC053 JC079 JC105 | JC001 vs JC027a | 0.965 |
| | | JC001 vs JC053 | 0.968 |
| | | JC001 vs JC079 | 0.96 |
| | | JC001 vs JC105 | 0.968 |
| | | JC027a vs JC105 | 0.952 |
| | | JC027a vs JC079 | 0.953 |
| | | JC027a vs JC053 | 0.962 |
| | | JC053 vs JC105 | 0.971 |
| | | JC053 vs JC079 | 0.985 |
| | | JC079 vs JC105 | 0.975 |
| g2 | JC002 JC028 JC054 JC080 JC106a | JC002 vs JC028 | 0.922 |
| | | JC002 vs JC054 | 0.922 |
| | | JC002 vs JC080 | 0.922 |
| | | JC002 vs JC106a | 0.928 |
| | | JC028 vs JC054 | 0.987 |
| | | JC028 vs JC080 | 0.98 |
| | | JC028 vs JC106a | 0.974 |
| | | JC054 vs JC080 | 0.984 |
| | | JC054 vs JC106a | 0.982 |
| | | JC080 vs JC106a | 0.966 |
| g3 | JC003 JC029a JC055 JC081 JC107a | JC003 vs JC029a | 0.989 |
| | | JC003 vs JC055 | 0.987 |
| | | JC003 vs JC081 | 0.954 |
| | | JC003 vs JC107a | 0.99 |
| | | JC029a vs JC055 | 0.984 |
| | | JC029a vs JC081 | 0.957 |
| | | JC029a vs JC107a | 0.985 |
| | | JC055 vs JC081 | 0.959 |
| | | JC055 vs JC107a | 0.983 |
| | | JC081 vs JC107a | 0.945 |
| g4 | JC004 JC030a JC056 JC082 | JC004 vs JC030a | 0.975 |
| | | JC004 vs JC056 | 0.976 |
| | | JC004 vs JC082 | 0.957 |
| | | JC004 vs JC108 | 0.945 |

| | | | |
|---|---|---|---|
| | JC108 | JC030a vs JC056 | 0.988 |
| | | JC030a vs JC082 | 0.976 |
| | | JC030a vs JC108 | 0.967 |
| | | JC056 vs JC082 | 0.983 |
| | | JC056 vs JC108 | 0.972 |
| | | JC082 vs JC108 | 0.984 |
| g5 | JC005<br>JC031<br>JC057<br>JC083a<br>JC109a | JC005 vs JC031 | 0.968 |
| | | JC005 vs JC057 | 0.98 |
| | | JC005 vs JC083a | 0.985 |
| | | JC005 vs JC109a | 0.985 |
| | | JC031 vs JC057 | 0.981 |
| | | JC031 vs JC083a | 0.963 |
| | | JC031 vs JC109a | 0.962 |
| | | JC057 vs JC083a | 0.975 |
| | | JC057 vs JC109a | 0.975 |
| | | JC083a vs JC109a | 0.991 |
| g6 | JC006<br>JC032<br>JC058<br>JC084<br>JC110a | JC006 vs JC032 | 0.978 |
| | | JC006 vs JC058 | 0.987 |
| | | JC006 vs JC084 | 0.97 |
| | | JC006 vs JC110a | 0.983 |
| | | JC032 vs JC058 | 0.969 |
| | | JC032 vs JC084 | 0.983 |
| | | JC032 vs JC110a | 0.959 |
| | | JC058 vs JC084 | 0.97 |
| | | JC058 vs JC110a | 0.982 |
| | | JC084 vs JC110a | 0.952 |
| g7 | JC007<br>JC033<br>JC059<br>JC085<br>JC111a | JC007 vs JC033 | 0.982 |
| | | JC007 vs JC059 | 0.988 |
| | | JC007 vs JC085 | 0.983 |
| | | JC007 vs JC111a | 0.988 |
| | | JC033 vs JC059 | 0.977 |
| | | JC033 vs JC085 | 0.96 |
| | | JC033 vs JC111a | 0.988 |
| | | JC059 vs JC085 | 0.984 |
| | | JC059 vs JC111a | 0.983 |
| | | JC085 vs JC111a | 0.973 |
| g8 | JC008<br>JC034a<br>JC060 | JC008 vs JC034a | 0.984 |
| | | JC008 vs JC060 | 0.986 |
| | | JC008 vs JC086 | 0.968 |

| | | | |
|---|---|---|---|
| | JC086<br>JC112 | JC008 vs JC112 | 0.979 |
| | | JC034a vs JC060 | 0.987 |
| | | JC034a vs JC086 | 0.967 |
| | | JC034a vs JC112 | 0.979 |
| | | JC060 vs JC086 | 0.964 |
| | | JC060 vs JC112 | 0.977 |
| | | JC086 vs JC112 | 0.984 |
| g9 | JC009J<br>C035a<br>JC061<br>JC087<br>JC113a | JC009 vs JC035a | 0.983 |
| | | JC009 vs JC061 | 0.981 |
| | | JC009 vs JC087 | 0.985 |
| | | JC009 vs JC113a | 0.983 |
| | | JC035a vs JC061 | 0.991 |
| | | JC035a vs JC087 | 0.974 |
| | | JC035a vs JC113a | 0.992 |
| | | JC061 vs JC087 | 0.97 |
| | | JC061 vs JC113a | 0.989 |
| | | JC087 vs JC113a | 0.974 |
| g10 | JC010<br>JC036a<br>JC062<br>JC088<br>JC114a | JC010 vs JC036a | 0.987 |
| | | JC010 vs JC062 | 0.986 |
| | | JC010 vs JC088 | 0.97 |
| | | JC010 vs JC114a | 0.987 |
| | | JC036a vs JC062 | 0.978 |
| | | JC036a vs JC088 | 0.966 |
| | | JC036a vs JC114a | 0.989 |
| | | JC062 vs JC088 | 0.972 |
| | | JC062 vs JC114a | 0.982 |
| | | JC088 vs JC114a | 0.969 |
| g11 | JC011<br>JC037a<br>JC063<br>JC089<br>JC115 | JC011 vs JC037a | 0.982 |
| | | JC011 vs JC063 | 0.985 |
| | | JC011 vs JC089 | 0.986 |
| | | JC011 vs JC115 | 0.986 |
| | | JC037a vs JC063 | 0.978 |
| | | JC037a vs JC089 | 0.982 |
| | | JC037a vs JC115 | 0.986 |
| | | JC063 vs JC089 | 0.984 |
| | | JC063 vs JC115 | 0.984 |
| | | JC089 vs JC115 | 0.987 |
| g12 | JC012<br>JC038a | JC012 vs JC038a | 0.969 |
| | | JC012 vs JC064 | 0.939 |

| | | | |
|---|---|---|---|
| | JC064<br>JC090<br>JC116 | JC012 vs JC090 | 0.964 |
| | | JC012 vs JC116 | 0.938 |
| | | JC038a vs JC064 | 0.974 |
| | | JC038a vs JC090 | 0.972 |
| | | JC038a vs JC116 | 0.952 |
| | | JC064 vs JC090 | 0.945 |
| | | JC064 vs JC116 | 0.933 |
| | | JC090 vs JC116 | 0.958 |
| g13 | JC013<br>JC039<br>JC065<br>JC091<br>JC117a | JC013 vs JC039 | 0.958 |
| | | JC013 vs JC065 | 0.982 |
| | | JC013 vs JC091 | 0.975 |
| | | JC013 vs JC117a | 0.978 |
| | | JC039 vs JC065 | 0.978 |
| | | JC039 vs JC091 | 0.97 |
| | | JC039 vs JC117a | 0.954 |
| | | JC065 vs JC091 | 0.985 |
| | | JC065 vs JC117a | 0.978 |
| | | JC091 vs JC117a | 0.973 |
| g14 | JC014<br>JC040a<br>JC066<br>JC092<br>JC118a | JC014 vs JC040a | 0.985 |
| | | JC014 vs JC066 | 0.991 |
| | | JC014 vs JC092 | 0.981 |
| | | JC014 vs JC118a | 0.988 |
| | | JC040a vs JC066 | 0.983 |
| | | JC040a vs JC092 | 0.968 |
| | | JC040a vs JC118a | 0.984 |
| | | JC066 vs JC092 | 0.984 |
| | | JC066 vs JC118a | 0.986 |
| | | JC092 vs JC118a | 0.984 |
| g15 | JC015<br>JC041a<br>JC067<br>JC093<br>JC119 | JC015 vs JC041a | 0.978 |
| | | JC015 vs JC067 | 0.98 |
| | | JC015 vs JC093 | 0.978 |
| | | JC015 vs JC119 | 0.975 |
| | | JC041a vs JC067 | 0.987 |
| | | JC041a vs JC093 | 0.983 |
| | | JC041a vs JC119 | 0.981 |
| | | JC067 vs JC093 | 0.991 |
| | | JC067 vs JC119 | 0.984 |
| | | JC093 vs JC119 | 0.981 |
| g16 | JC016 | JC016 vs JC042 | 0.979 |

| | | | |
|---|---|---|---|
| | JC042<br>JC068<br>JC094<br>JC120 | JC016 vs JC068 | 0.974 |
| | | JC016 vs JC094 | 0.974 |
| | | JC016 vs JC120 | 0.978 |
| | | JC042 vs JC068 | 0.967 |
| | | JC042 vs JC094 | 0.962 |
| | | JC042 vs JC120 | 0.982 |
| | | JC068 vs JC094 | 0.967 |
| | | JC068 vs JC120 | 0.969 |
| | | JC094 vs JC120 | 0.964 |
| g17 | JC017<br>JC043a<br>JC069<br>JC095<br>JC121 | JC017 vs JC043a | 0.981 |
| | | JC017 vs JC069 | 0.978 |
| | | JC017 vs JC095 | 0.969 |
| | | JC017 vs JC121 | 0.972 |
| | | JC043a vs JC069 | 0.983 |
| | | JC043a vs JC095 | 0.973 |
| | | JC043a vs JC121 | 0.973 |
| | | JC069 vs JC095 | 0.978 |
| | | JC069 vs JC121 | 0.977 |
| | | JC095 vs JC121 | 0.98 |
| g18 | JC018<br>JC044a<br>JC070<br>JC096<br>JC122 | JC018 vs JC044a | 0.985 |
| | | JC018 vs JC070 | 0.985 |
| | | JC018 vs JC096 | 0.955 |
| | | JC018 vs JC122 | 0.983 |
| | | JC044a vs JC070 | 0.98 |
| | | JC044a vs JC096 | 0.952 |
| | | JC044a vs JC122 | 0.981 |
| | | JC070 vs JC096 | 0.966 |
| | | JC070 vs JC122 | 0.987 |
| | | JC096 vs JC122 | 0.959 |
| g19 | JC019<br>JC045a<br>JC071<br>JC097<br>JC123a | JC019 vs JC045a | 0.974 |
| | | JC019 vs JC097 | 0.982 |
| | | JC019 vs JC123a | 0.983 |
| | | JC045a vs JC071 | 0.96 |
| | | JC045a vs JC123a | 0.985 |
| | | JC097 vs JC123a | 0.977 |
| | | JC071 vs JC097 | 0.978 |
| g20 | JC020<br>JC046a<br>JC072 | JC020 vs JC046a | 0.969 |
| | | JC020 vs JC072 | 0.986 |
| | | JC020 vs JC098 | 0.979 |

| | | | |
|---|---|---|---|
| | JC098 JC124a | JC020 vs JC124a | 0.979 |
| | | JC046a vs JC072 | 0.965 |
| | | JC046a vs JC098 | 0.969 |
| | | JC046a vs JC124a | 0.985 |
| | | JC072 vs JC098 | 0.98 |
| | | JC072 vs JC124a | 0.959 |
| | | JC098 vs JC124a | 0.983 |
| g21 | JC021 JC047a JC073 JC099a JC125a | JC021 vs JC047a | 0.979 |
| | | JC021 vs JC073 | 0.977 |
| | | JC021 vs JC099a | 0.973 |
| | | JC021 vs JC125a | 0.974 |
| | | JC047a vs JC073 | 0.968 |
| | | JC047a vs JC099a | 0.991 |
| | | JC047a vs JC125a | 0.973 |
| | | JC073 vs JC099a | 0.959 |
| | | JC073 vs JC125a | 0.971 |
| | | JC099a vs JC125a | 0.969 |
| g22 | JC022 JC048a JC074 JC100 JC126 | JC022 vs JC048a | 0.976 |
| | | JC022 vs JC074 | 0.989 |
| | | JC022 vs JC100 | 0.97 |
| | | JC022 vs JC126 | 0.983 |
| | | JC048a vs JC074 | 0.975 |
| | | JC048a vs JC100 | 0.96 |
| | | JC048a vs JC126 | 0.969 |
| | | JC074 vs JC100 | 0.979 |
| | | JC074 vs JC126 | 0.988 |
| | | JC100 vs JC126 | 0.984 |
| g23 | JC023 JC049a JC075 JC101 JC127 | JC023 vs JC049a | 0.943 |
| | | JC023 vs JC075 | 0.951 |
| | | JC023 vs JC101 | 0.96 |
| | | JC023 vs JC127 | 0.955 |
| | | JC049a vs JC075 | 0.981 |
| | | JC049a vs JC101 | 0.97 |
| | | JC049a vs JC127 | 0.972 |
| | | JC075 vs JC101 | 0.984 |
| | | JC075 vs JC127 | 0.987 |
| | | JC101 vs JC127 | 0.986 |
| g24 | JC024 JC050a | JC024 vs JC050a | 0.969 |
| | | JC024 vs JC076 | 0.979 |

| | | | |
|---|---|---|---|
| | JC076<br>JC128 | JC024 vs JC028 | 0.982 |
| | | JC050a vs JC076 | 0.979 |
| | | JC050a vs JC028 | 0.969 |
| | | JC076 vs JC028 | 0.98 |
| g25 | JC025<br>JC051a<br>JC077<br>JC103<br>JC129 | JC025 vs JC051a | 0.972 |
| | | JC025 vs JC077 | 0.984 |
| | | JC025 vs JC103 | 0.981 |
| | | JC025 vs JC129 | 0.985 |
| | | JC051a vs JC077 | 0.985 |
| | | JC051a vs JC103 | 0.98 |
| | | JC051a vs JC129 | 0.971 |
| | | JC077 vs JC103 | 0.988 |
| | | JC077 vs JC129 | 0.982 |
| | | JC103 vs JC129 | 0.985 |
| g26 | JC026<br>JC052a<br>JC078<br>JC130 | JC026 vs JC052a | 0.982 |
| | | JC026 vs JC078 | 0.987 |
| | | JC026 vs JC130 | 0.979 |
| | | JC052a vs JC078 | 0.988 |
| | | JC052a vs JC130 | 0.973 |
| | | JC078 vs JC130 | 0.977 |

**S3 Table Genes uniquely expressed in two strains of mice under different conditions participating in metabolic processes.**

| Gene | Control | | Glucose | | Sucrose | | Fructose | |
|---|---|---|---|---|---|---|---|---|
| | C57BL/6J | BALB/cJ | C57BL/6J | BALB/cJ | C57BL/6J | BALB/cJ | C57BL/6J | BALB/cJ |
| | in g11only | in g15only | in g12only | in g16only | in g13only | in g17only | in g14only | in g18only |
| *Oasl2* | *Oasl2* | - | *Oasl2* | - | *Oasl2* | - | - | - |
| *Srd5a2* | *Srd5a2* | - | *Srd5a2* | - | - | - | - | *Srd5a2* |
| *Pfkfb3* | *Pfkfb3* | - | - | - | - | - | - | - |
| *Akr1cl* | *Akr1cl* | - | - | - | - | - | - | - |
| *Ankrd2* | *Ankrd2* | - | *Ankrd2* | - | *Ankrd2* | - | *Ankrd2* | - |
| *Anxa9* | *Anxa9* | - | - | - | - | - | - | - |
| *Asns* | *Asns* | - | *Asns* | - | *Asns* | - | *Asns* | - |
| *Batf3* | *Batf3* | - | *Batf3* | - | - | - | *Batf3* | - |
| *Capn8* | *Capn8* | - | *Capn8* | - | *Capn8* | - | *Capn8* | - |
| *Ccl5* | *Ccl5* | - | - | - | *Ccl5* | - | *Ccl5* | - |
| *Cyp11a1* | *Cyp11a1* | - | *Cyp11a1* | - | - | - | - | *Cyp11a1* |
| *Cela3b* | *Cela3b* | - | *Cela3b* | - | - | - | *Cela3b* | - |
| *Ten1* | *Ten1* | - | - | *Ten1* | *Ten1* | - | - | - |
| *Cdkl2* | *Cdkl2* | - | *Cdkl2* | - | - | - | - | - |
| *Cybb* | *Cybb* | - | *Cybb* | - | - | *Cybb* | *Cybb* | - |
| *Cyp11b1* | *Cyp11b1* | - | *Cyp11b1* | - | - | - | - | *Cyp11b1* |
| *Cyp26c1* | *Cyp26c1* | - | *Cyp26c1* | - | *Cyp26c1* | - | *Cyp26c1* | - |
| *Cyp2c69* | *Cyp2c69* | - | *Cyp2c69* | - | *Cyp2c69* | - | *Cyp2c69* | - |
| *Ube3d* | *Ube3d* | - | *Ube3d* | - | - | - | *Ube3d* | - |
| *Egr1* | *Egr1* | - | - | - | - | - | - | - |
| *Art4* | *Art4* | - | *Art4* | - | *Art4* | - | *Art4* | - |
| *Fancc* | *Fancc* | - | *Fancc* | - | *Fancc* | - | *Fancc* | - |

| | | | | | | | | |
|---|---|---|---|---|---|---|---|---|
| *Fbxl21* | *Fbxl21* | - | *Fbxl21* | - | *Fbxl21* | - | *Fbxl21* | - |
| *Fdxacb1* | *Fdxacb1* | - | - | - | - | - | - | - |
| *Fosl2* | *Fosl2* | - | - | - | - | - | - | - |
| *Gmds* | *Gmds* | - | *Gmds* | - | *Gmds* | - | *Gmds* | - |
| *Serpine2* | *Serpine2* | - | *Serpine2* | - | *Serpine2* | - | *Serpine2* | - |
| *Gpat2* | *Gpat2* | - | *Gpat2* | - | *Gpat2* | - | *Gpat2* | - |
| *Tdg* | *Tdg* | - | *Tdg* | - | *Tdg* | - | *Tdg* | - |
| *Hist1h2ab* | *Hist1h2ab* | - | - | - | *Hist1h2ab* | - | - | - |
| *Hist1h2ac* | *Hist1h2ac* | - | - | - | *Hist1h2ac* | - | - | - |
| *Hist1h2ad* | *Hist1h2ad* | - | - | - | *Hist1h2ad* | - | - | - |
| *Hist1h2ae* | *Hist1h2ae* | - | - | - | *Hist1h2ae* | - | - | - |
| *Hist1h2ag* | *Hist1h2ag* | - | - | - | *Hist1h2ag* | - | - | - |
| *Hist1h2ai* | *Hist1h2ai* | - | - | - | *Hist1h2ai* | - | - | - |
| *Hist1h2an* | *Hist1h2an* | - | - | - | *Hist1h2an* | - | - | - |
| *Hist1h2ao* | *Hist1h2ao* | - | - | - | *Hist1h2ao* | - | - | - |
| *Hist1h2ap* | *Hist1h2ap* | - | - | - | *Hist1h2ap* | - | - | - |
| *Hist3h2a* | *Hist3h2a* | - | - | - | *Hist3h2a* | - | *Hist3h2a* | - |
| *Smyd1* | *Smyd1* | - | *Smyd1* | - | - | - | *Smyd1* | - |
| *Inhba* | *Inhba* | - | - | - | - | - | - | - |
| *Ints12* | *Ints12* | - | - | - | - | - | - | - |
| *Klf11* | *Klf11* | - | - | - | - | - | - | - |
| *Dct* | *Dct* | - | *Dct* | - | *Dct* | - | - | - |
| *Serpinb1a* | *Serpinb1a* | - | *Serpinb1a* | - | - | - | *Serpinb1a* | - |
| *Ajuba* | *Ajuba* | - | *Ajuba* | - | *Ajuba* | - | *Ajuba* | - |
| *Mapkapk3* | *Mapkapk3* | - | *Mapkapk3* | - | *Mapkapk3* | - | *Mapkapk3* | - |
| *B3gnt3* | *B3gnt3* | - | *B3gnt3* | - | *B3gnt3* | - | - | - |
| *Ncf1* | *Ncf1* | - | *Ncf1* | - | - | - | *Ncf1* | - |
| *Pck2* | *Pck2* | - | *Pck2* | - | *Pck2* | - | *Pck2* | - |
| *Pms1* | *Pms1* | - | - | - | - | - | - | - |

| | | | | | | | | |
|---|---|---|---|---|---|---|---|---|
| *Eme2* | *Eme2* | - | - | - | *Eme2* | - | *Eme2* | - |
| *Zdhhc21* | *Zdhhc21* | - | *Zdhhc21* | - | - | - | - | - |
| *Rrm2* | *Rrm2* | - | *Rrm2* | - | *Rrm2* | - | *Rrm2* | - |
| *Rik* | *Rik* | - | *Rik* | - | *Rik* | - | *Rik* | - |
| *Spag8* | *Spag8* | - | - | *Spag8* | - | - | - | - |
| *Socs2* | *Socs2* | - | - | - | *Socs2* | - | - | - |
| *Traf3ip2* | *Traf3ip2* | - | *Traf3ip2* | - | - | - | - | - |
| *E2f1* | *E2f1* | - | - | - | *E2f1* | - | *E2f1* | - |
| *Hck* | *Hck* | - | *Hck* | - | - | - | *Hck* | - |
| *Zap70* | *Zap70* | - | *Zap70* | - | *Zap70* | - | - | - |
| *Ube2u* | *Ube2u* | - | *Ube2u* | - | *Ube2u* | - | *Ube2u* | - |
| *Ugt1a5* | *Ugt1a5* | - | *Ugt1a5* | - | - | - | *Ugt1a5* | - |
| *Zfp113* | *Zfp113* | - | - | - | - | - | - | - |
| *Znf296* | *Znf296* | - | - | *Znf296* | *Znf296* | - | - | - |
| *Zhx2* | *Zhx2* | - | *Zhx2* | - | *Zhx2* | - | | - |
| *Primpol* | - | *Primpol* | - | - | - | - | - | *Primpol* |
| *Ptprs* | - | *Ptprs* | - | - | - | - | - | - |
| *AK157302* | - | *AK157302* | - | *AK157302* | - | *AK157302* | - | *AK157302* |
| *Slx4* | - | *Slx4* | - | *Slx4* | - | - | - | *Slx4* |
| *Plau* | - | *Plau* | - | *Plau* | - | - | - | - |
| *Gm5294* | - | *Gm5294* | - | *Gm5294* | - | *Gm5294* | - | *Gm5294* |
| *Ncf4* | - | *Ncf4* | - | - | - | - | - | - |
| *P3h3* | - | *P3h3* | - | - | - | - | - | *P3h3* |
| *Otub2* | - | *Otub2* | - | - | - | - | - | - |
| *Wars2* | - | *Wars2* | - | - | - | - | - | - |
| *Cdk19* | - | *Cdk19* | - | - | - | - | - | - |
| *Il1a* | - | *Il1a* | - | - | - | - | - | - |
| *Hmga1* | - | *Hmga1* | - | - | - | *Hmga1* | - | *Hmga1* |
| *Gm10639* | - | *Gm10639* | - | *Gm10639* | - | *Gm10639* | - | *Gm10639* |

| | | | | | | | | |
|---|---|---|---|---|---|---|---|---|
| *Camta1* | - | *Camta1* | - | - | - | - | - | - |
| *Mmp2* | - | *Mmp2* | - | *Mmp2* | - | *Mmp2* | - | *Mmp2* |
| *Gm6665* | - | *Gm6665* | - | *Gm6665* | - | - | - | *Gm6665* |
| *Mcm4* | - | *Mcm4* | - | *Mcm4* | - | - | - | *Mcm4* |
| *E2f7* | - | *E2f7* | - | *E2f7* | - | *E2f7* | - | *E2f7* |
| *Pus7l* | - | *Pus7l* | - | - | - | - | - | - |
| *Capn3* | - | *Capn3* | - | *Capn3* | - | *Capn3* | - | *Capn3* |
| *Mycb* | - | *Mycb* | - | *Mycb* | - | *Mycb* | - | *Mycb* |
| *Gls* | - | *Gls* | - | - | - | - | - | - |
| *Mcm2* | - | *Mcm2* | - | *Mcm2* | - | - | - | *Mcm2* |
| *Rev3l* | - | *Rev3l* | - | - | - | *Rev3l* | - | - |
| *Iglc2* | - | *Iglc2* | - | *Iglc2* | - | *Iglc2* | - | *Iglc2* |
| *Cyp2g1* | - | *Cyp2g1* | - | - | - | *Cyp2g1* | - | - |
| *Igha* | - | *Igha* | *Igha* | - | - | - | - | - |
| *Miox* | - | *Miox* | - | *Miox* | - | - | - | - |
| *Pou6f1* | - | *Pou6f1* | - | - | - | - | - | - |
| *Fan1* | - | *Fan1* | - | - | - | *Fan1* | - | - |
| *Exog* | - | *Exog* | - | *Exog* | - | - | - | - |
| *Gm12248* | - | *Gm12248* | - | *Gm12248* | - | *Gm12248* | - | *Gm12248* |
| *Trmt44* | - | *Trmt44* | - | - | - | - | - | - |
| *Mast1* | - | *Mast1* | - | *Mast1* | - | *Mast1* | - | *Mast1* |
| *Sgsh* | - | *Sgsh* | - | *Sgsh* | - | - | - | - |
| *Usp37* | - | *Usp37* | - | - | - | *Usp37* | - | *Usp37* |
| *Il1b* | - | *Il1b* | - | - | - | - | - | - |
| *Src* | - | *Src* | - | - | - | - | - | - |
| *Pgm5* | - | *Pgm5* | - | *Pgm5* | - | *Pgm5* | - | - |
| *Mzb1* | - | *Mzb1* | - | *Mzb1* | - | - | - | - |
| *Mettl15* | - | *Mettl15* | - | - | - | - | - | - |
| *Nhlrc1* | - | *Nhlrc1* | - | *Nhlrc1* | - | *Nhlrc1* | - | - |

| | | | | | | | | |
|---|---|---|---|---|---|---|---|---|
| *Tp53bp1* | - | *Tp53bp1* | - | - | - | *Tp53bp1* | - | *Tp53bp1* |
| *Gspt2* | - | *Gspt2* | - | *Gspt2* | - | *Gspt2* | - | *Gspt2* |
| *Zfp142* | - | *Zfp142* | - | - | - | - | - | *Zfp142* |
| *Lhx6* | - | *Lhx6* | - | - | - | - | - | - |
| *Pars2* | - | *Pars2* | - | *Pars2* | - | *Pars2* | - | *Pars2* |
| *Nupr2* | - | *Nupr2* | - | - | - | *Nupr2* | - | - |
| *Gucy2c* | - | *Gucy2c* | - | *Gucy2c* | - | *Gucy2c* | - | *Gucy2c* |
| *Cyfip2* | - | *Cyfip2* | - | *Cyfip2* | - | *Cyfip2* | - | *Cyfip2* |
| *Dtx2* | - | *Dtx2* | - | - | - | - | - | - |
| *Elk1* | - | *Elk1* | - | - | - | - | - | - |
| *Recql* | - | *Recql* | - | - | - | - | - | - |
| *Gdpd1* | - | *Gdpd1* | - | - | - | *Gdpd1* | - | *Gdpd1* |
| *Mt3* | - | *Mt3* | - | *Mt3* | - | - | - | - |
| *Gm9774* | - | *Gm9774* | - | *Gm9774* | - | - | - | *Gm9774* |
| *Fli1* | - | *Fli1* | - | - | - | - | - | - |
| *BC005561* | - | *BC005561* | - | - | - | - | - | - |
| *Cyp2a4* | - | *Cyp2a4* | - | - | - | - | - | *Cyp2a4* |
| *Spindoc* | - | *Spindoc* | - | *Spindoc* | - | - | - | - |
| *Serpina3c* | - | *Serpina3c* | - | *Serpina3c* | - | *Serpina3c* | - | *Serpina3c* |
| *Akr1c18* | - | *Akr1c18* | - | *Akr1c18* | - | - | - | - |
| *Ggt1* | - | *Ggt1* | - | *Ggt1* | - | - | - | - |
| *Suv39h1* | - | *Suv39h1* | - | - | *Suv39h1* | - | - | *Suv39h1* |
| *Rad9b* | - | *Rad9b* | - | *Rad9b* | - | *Rad9b* | - | *Rad9b* |
| *Ccnj* | - | *Ccnj* | - | *Ccnj* | - | - | - | - |
| *Cers5* | - | *Cers5* | - | - | - | - | - | - |
| *Klf12* | - | *Klf12* | - | - | - | *Klf12* | - | *Klf12* |
| *Elf5* | - | *Elf5* | - | *Elf5* | - | *Elf5* | - | *Elf5* |
| *Acsm2* | - | *Acsm2* | - | *Acsm2* | - | - | - | - |
| *Mcm3* | - | *Mcm3* | - | *Mcm3* | - | - | - | *Mcm3* |

| | | | | | | | | |
|---|---|---|---|---|---|---|---|---|
| *Ppargc1a* | - | *Ppargc1a* | - | - | - | *Ppargc1a* | - | - |
| *Pgap3* | - | *Pgap3* | - | *Pgap3* | - | - | - | *Pgap3* |
| *Fbxo17* | - | *Fbxo17* | - | *Fbxo17* | - | *Fbxo17* | - | *Fbxo17* |
| *Neu3* | - | *Neu3* | - | - | - | *Neu3* | - | - |
| *Kctd10* | - | *Kctd10* | - | *Kctd10* | - | *Kctd10* | - | - |
| *Far1* | - | *Far1* | - | - | - | - | - | - |
| *Serpina5* | - | *Serpina5* | - | *Serpina5* | - | *Serpina5* | - | *Serpina5* |
| *Crtc1* | - | *Crtc1* | - | - | - | - | - | - |
| *Prkdc* | - | *Prkdc* | - | *Prkdc* | - | *Prkdc* | - | *Prkdc* |
| *Bora* | - | *Bora* | - | *Bora* | - | - | - | *Bora* |
| *Timp3* | - | *Timp3* | - | - | - | - | *Timp3* | - |
| *Serpina9* | - | *Serpina9* | - | - | - | - | - | - |
| *Depdc5* | - | *Depdc5* | - | - | - | - | - | - |
| *Nup133* | - | *Nup133* | - | - | - | - | - | - |
| *Tbx2* | - | *Tbx2* | - | - | - | - | - | - |
| *Serpina3a* | - | *Serpina3a* | - | *Serpina3a* | - | *Serpina3a* | - | *Serpina3a* |
| *Carf* | - | *Carf* | - | - | - | - | - | - |
| *Helz* | - | *Helz* | - | - | - | - | - | - |
| *Lhx2* | - | *Lhx2* | - | *Lhx2* | - | *Lhx2* | - | *Lhx2* |
| *Dcp1b* | - | *Dcp1b* | - | *Dcp1b* | - | - | - | - |
| *Synj2* | - | *Synj2* | - | - | - | *Synj2* | - | *Synj2* |
| *Tbp* | - | *Tbp* | - | - | - | *Tbp* | - | - |
| *Tarbp1* | - | *Tarbp1* | - | *Tarbp1* | - | *Tarbp1* | - | *Tarbp1* |
| *Sufu* | - | *Sufu* | - | *Sufu* | - | - | - | - |
| *Pnlip* | - | - | *Pnlip* | - | - | - | *Pnlip* | - |
| *Cpa2* | - | - | *Cpa2* | - | - | - | - | - |
| *Atad2b* | - | - | *Atad2b* | - | - | - | *Atad2b* | - |
| *Reck* | - | - | *Reck* | - | *Reck* | - | *Reck* | - |
| *Rik* | - | - | *Rik* | - | - | - | - | - |

| | | | | | | | | |
|---|---|---|---|---|---|---|---|---|
| *Chil3* | - | - | *Chil3* | - | - | - | - | - |
| *Zfp407* | - | - | *Zfp407* | - | - | - | - | - |
| *Hk3* | - | - | *Hk3* | - | - | - | - | - |
| *Apobec3* | - | - | *Apobec3* | - | *Apobec3* | - | *Apobec3* | - |
| *Cela2a* | - | - | *Cela2a* | - | - | - | - | - |
| *Map3k21* | - | - | *Map3k21* | - | *Map3k21* | - | - | - |
| *Polk* | - | - | *Polk* | - | *Polk* | - | *Polk* | - |
| *Pot1b* | - | - | *Pot1b* | - | *Pot1b* | - | *Pot1b* | - |
| *Pgap1* | - | - | *Pgap1* | - | *Pgap1* | - | *Pgap1* | - |
| *Prss2* | - | - | *Prss2* | - | - | - | *Prss2* | - |
| *Cpb1* | - | - | *Cpb1* | - | - | - | - | - |
| *Tmem173* | - | - | *Tmem173* | - | - | *Tmem173* | - | - |
| *Fam126a* | - | - | *Fam126a* | - | - | - | *Fam126a* | - |
| *Etaa1* | - | - | *Etaa1* | - | - | - | - | - |
| *Prkch* | - | - | *Prkch* | - | - | - | - | - |
| *Ptges* | - | - | *Ptges* | - | *Ptges* | - | *Ptges* | - |
| *Serpinb6b* | - | - | *Serpinb6b* | - | *Serpinb6b* | - | *Serpinb6b* | - |
| *Oas1a* | - | - | *Oas1a* | - | - | - | *Oas1a* | - |
| *Relb* | - | - | *Relb* | - | - | - | - | - |
| *Il1rn* | - | - | *Il1rn* | - | - | - | - | - |
| *Try4* | - | - | *Try4* | - | - | - | *Try4* | - |
| *Hip1* | - | - | *Hip1* | - | - | - | - | - |
| *Pik3c2b* | - | - | *Pik3c2b* | - | *Pik3c2b* | - | *Pik3c2b* | - |
| *Capn11* | - | - | *Capn11* | - | - | - | - | - |
| *Rnf152* | - | - | *Rnf152* | - | - | - | - | - |
| *Map3k13* | - | - | *Map3k13* | - | - | - | *Map3k13* | - |
| *Try5* | - | - | *Try5* | - | - | *Try5* | - | - |
| *Pnliprp2* | - | - | *Pnliprp2* | - | - | - | - | - |
| *Fgr* | - | - | *Fgr* | - | - | - | *Fgr* | - |

| | | | | | | | | | |
|---|---|---|---|---|---|---|---|---|---|
| *Pdgfc* | - | - | *Pdgfc* | - | - | - | *Pdgfc* | - | - |
| *Ccl2* | - | - | *Ccl2* | - | *Ccl2* | - | *Ccl2* | - |
| *Serpina3f* | - | - | *Serpina3f* | - | - | - | - | - |
| *Cpa1* | - | - | *Cpa1* | - | - | - | *Cpa1* | - |
| *Kit* | - | - | *Kit* | - | - | - | - | - |
| *Pm20d2* | - | - | *Pm20d2* | - | - | - | *Pm20d2* | - |
| *Pnliprp1* | - | - | *Pnliprp1* | - | - | - | - | - |
| *Bmp2k* | - | - | *Bmp2k* | - | - | - | *Bmp2k* | - |
| *Hist1h1d* | - | - | - | *Hist1h1d* | *Hist1h1d* | - | - | - |
| *Rpusd1* | - | - | - | *Rpusd1* | - | - | - | - |
| *Tcf7* | - | - | - | *Tcf7* | - | *Tcf7* | - | - |
| *Gdpgp1* | - | - | - | *Gdpgp1* | - | - | - | - |
| *Apobec2* | - | - | - | *Apobec2* | - | - | - | - |
| *Azin2* | - | - | - | *Azin2* | - | *Azin2* | - | *Azin2* |
| *Thada* | - | - | - | *Thada* | - | - | - | - |
| *Ppargc1b* | - | - | - | *Ppargc1b* | - | *Ppargc1b* | - | - |
| *Glis2* | - | - | - | *Glis2* | - | - | - | - |
| *Hist1h1e* | - | - | - | *Hist1h1e* | - | - | - | - |
| *Aph1b* | - | - | - | *Aph1b* | - | - | - | - |
| *Pfkfb4* | - | - | - | *Pfkfb4* | - | *Pfkfb4* | - | - |
| *Sox12* | - | - | - | *Sox12* | - | - | *Sox12* | - |
| *Fut8* | - | - | - | *Fut8* | - | *Fut8* | - | - |
| *Pde6g* | - | - | - | *Pde6g* | - | - | - | - |
| *Rad51c* | - | - | - | *Rad51c* | - | - | - | *Rad51c* |
| *Akr1c21* | - | - | - | *Akr1c21* | - | - | - | - |
| *Dnase1* | - | - | - | *Dnase1* | - | - | - | - |
| *Pld6* | - | - | - | *Pld6* | - | *Pld6* | - | *Pld6* |
| *Nr1h5* | - | - | - | *Nr1h5* | - | - | - | - |
| *Ripply3* | - | - | - | *Ripply3* | - | - | - | - |

| | | | | | | | | |
|---|---|---|---|---|---|---|---|---|
| *Gm4450* | - | - | - | *Gm4450* | - | - | - | - |
| *Hk1* | - | - | - | *Hk1* | - | - | - | - |
| *Ctsk* | - | - | - | *Ctsk* | - | - | - | - |
| *Rfx5* | - | - | - | *Rfx5* | - | - | - | - |
| *Tfcp2l1* | - | - | - | *Tfcp2l1* | - | - | - | - |
| *Aebp1* | - | - | - | *Aebp1* | - | - | - | - |
| *Timp1* | - | - | - | *Timp1* | - | - | - | - |
| *Hdac8* | - | - | - | *Hdac8* | - | *Hdac8* | - | - |
| *Irak1bp1* | - | - | - | *Irak1bp1* | - | *Irak1bp1* | - | - |
| *Rpusd2* | - | - | - | *Rpusd2* | - | - | - | - |
| *E2f2* | - | - | - | *E2f2* | - | *E2f2* | - | *E2f2* |
| *Fst* | - | - | - | - | *Fst* | - | - | - |
| *Cyp2d11* | - | - | - | - | *Cyp2d11* | - | *Cyp2d11* | - |
| *Ihh* | - | - | - | - | *Ihh* | - | - | - |
| *Ces2c* | - | - | - | - | *Ces2c* | - | *Ces2c* | - |
| *Pagr1a* | - | - | - | - | *Pagr1a* | - | - | *Pagr1a* |
| *Cish* | - | - | - | - | *Cish* | - | - | *Cish* |
| *Trbc1* | - | - | - | - | *Trbc1* | - | *Trbc1* | - |
| *Rpl24* | - | - | - | - | *Rpl24* | - | - | - |
| *Rec8* | - | - | - | - | *Rec8* | - | *Rec8* | - |
| *Trbc2* | - | - | - | - | *Trbc2* | - | - | - |
| *Ighv1-56* | - | - | - | - | *Ighv1-56* | - | *Ighv1-56* | - |
| *Lat* | - | - | - | - | *Lat* | - | - | - |
| *Fam126b* | - | - | - | - | *Fam126b* | - | *Fam126b* | - |
| *Art3* | - | - | - | - | *Art3* | - | *Art3* | - |
| *Usp35* | - | - | - | - | *Usp35* | - | *Usp35* | - |
| *Cmpk2* | - | - | - | - | *Cmpk2* | - | *Cmpk2* | - |
| *Dnd1* | - | - | - | - | *Dnd1* | - | - | - |
| *Bmp6* | - | - | - | - | *Bmp6* | - | *Bmp6* | - |

| | | | | | | | | |
|---|---|---|---|---|---|---|---|---|
| *Wnk4* | - | - | - | - | *Wnk4* | - | *Wnk4* | - |
| *Ovgp1* | - | - | - | - | *Ovgp1* | - | - | - |
| *Tbx6* | - | - | - | - | *Tbx6* | - | - | *Tbx6* |
| *Ckmt1* | - | - | - | - | *Ckmt1* | - | - | - |
| *Ccnd1* | - | - | - | - | *Ccnd1* | - | *Ccnd1* | - |
| *Cyp46a1* | - | - | - | - | *Cyp46a1* | - | *Cyp46a1* | - |
| *Pkn3* | - | - | - | - | *Pkn3* | - | *Pkn3* | - |
| *Eif4ebp3* | - | - | - | - | - | *Eif4ebp3* | - | - |
| *Prkx* | - | - | - | - | - | *Prkx* | *Prkx* | - |
| *Dclre1c* | - | - | - | - | - | *Dclre1c* | - | - |
| *Znrf3* | - | - | - | - | - | *Znrf3* | *Znrf3* | - |
| *Anapc10* | - | - | - | - | - | *Anapc10* | - | - |
| *Mterf2* | - | - | - | - | - | *Mterf2* | - | *Mterf2* |
| *Spic* | - | - | - | - | - | *Spic* | - | - |
| *Cyp2c39* | - | - | - | - | - | *Cyp2c39* | - | *Cyp2c39* |
| *Lyl1* | - | - | - | - | - | *Lyl1* | - | - |
| *Cyp2b9* | - | - | - | - | - | *Cyp2b9* | - | *Cyp2b9* |
| *Lin54* | - | - | - | - | - | *Lin54* | - | *Lin54* |
| *Tspan15* | - | - | - | - | - | *Tspan15* | - | - |
| *Fuca2* | - | - | - | - | - | *Fuca2* | - | - |
| *Uba6* | - | - | - | - | - | - | *Uba6* | - |
| *Bbx* | - | - | - | - | - | - | *Bbx* | - |
| *Tnks* | - | - | - | - | - | - | *Tnks* | - |
| *Polh* | - | - | - | - | - | - | *Polh* | - |
| *Casp4* | - | - | - | - | - | - | *Casp4* | - |
| *Sall2* | - | - | - | - | - | - | *Sall2* | - |
| *Ints6l* | - | - | - | - | - | - | *Ints6l* | - |
| *Cyp2c55* | - | - | - | - | - | - | *Cyp2c55* | - |
| *Atf3* | - | - | - | - | - | - | *Atf3* | - |

| | | | | | | | | |
|---|---|---|---|---|---|---|---|---|
| *Rsf1* | - | - | - | - | - | - | *Rsf1* | - |
| *Serpinb9* | - | - | - | - | - | - | *Serpinb9* | - |
| *Nkx2-6* | - | - | - | - | - | - | *Nkx2-6* | - |
| *Celf2* | - | - | - | - | - | - | *Celf2* | - |
| *Gemin8* | - | - | - | - | - | - | - | *Gemin8* |
| *Etv5* | - | - | - | - | - | - | - | *Etv5* |
| *Ggt5* | - | - | - | - | - | - | - | *Ggt5* |
| *Th* | - | - | - | - | - | - | - | *Th* |
| *Aph1c* | - | - | - | - | - | - | - | *Aph1c* |
| *Dbh* | - | - | - | - | - | - | - | *Dbh* |

Note: uniquely expressed genes in either C57BL/6J or BALB/cJ mice at 6 hour postprandial fed without or with supplement of 20% of glucose, sucrose, or fructose in drinking water shown in Fig. 2 were annotated and classified with GO terms using Panther Classification system. Genes participating in metabolic processes are listed.

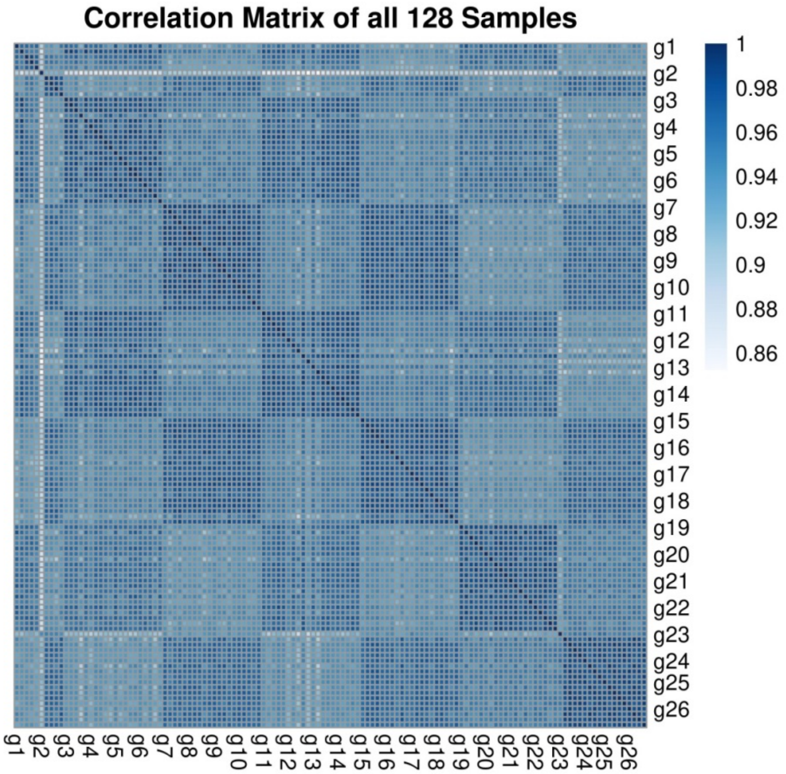

**S1 Fig. Correlation of gene expressions**. Correlation matrix of all 128 samples in 26 groups.

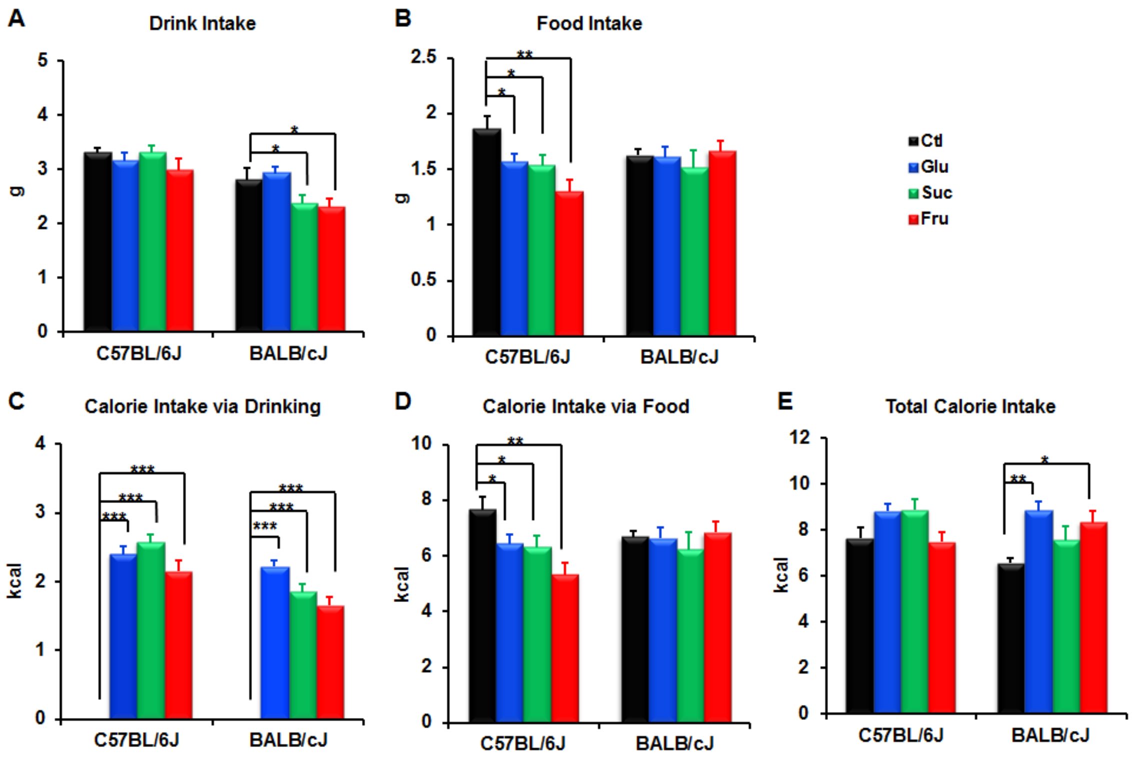

**S2 Fig. Drink, food and calorie intake.** Drink intake (A), food intake (B), and calorie intake via drinking (C) or food (D), or total calorie intake (C) of mice during the first 3 hours when 20% of glucose, sucrose, or fructose was separately supplemented in drinking water, n = 7-9 per group. Value are mean ± S.D. p values were obtained by two tail student t-test. *$p < 0.05$, **$p < 0.01$, or ***$p < 0.001$ was considered significant. Unit calories: glucose, 3.8 kcal/g; sucrose, 3.9 kcal/g; fructose 3.6 kcal/g; food, 4.11 kcal/g.
